# Stress-induced translation inhibition through rapid displacement of scanning initiation factors

**DOI:** 10.1101/2020.05.14.096354

**Authors:** Stefan Bresson, Vadim Shchepachev, Christos Spanos, Tomasz Turowski, Juri Rappsilber, David Tollervey

## Abstract

Cellular responses to environmental stress are frequently mediated by RNA-binding proteins (RBPs). Here, we examined global RBP dynamics in *Saccharomyces cerevisiae* in response to glucose starvation and heat shock. Each stress induced rapid remodeling of the RNA-protein interactome, without corresponding changes in RBP abundance. Consistent with general translation shutdown, ribosomal proteins contacting the mRNA showed decreased RNA-association. Among translation components, RNA-association was most reduced for initiation factors involved in 40S scanning (eIF4A, eIF4B, and Ded1), indicating a common mechanism of translational repression. In unstressed cells, eIF4A, eIF4B, and Ded1 primarily targeted the 5′-ends of mRNAs. Following glucose withdrawal, 5’-binding was abolished within 30sec, explaining the rapid translation shutdown, but mRNAs remained stable. Heat shock induced progressive loss of 5’ RNA-binding by initiation factors over ∼16min. Translation shutoff provoked selective 5′-degradation of mRNAs encoding translation-related factors, mediated by Xrn1. These results reveal mechanisms underlying translational control of gene expression during stress.

**Highlights:** A quantitative proteomic approach reveals rapid stress-induced changes in RNA-binding Translation shutdown is driven by loss of mRNA binding by scanning initiation factors eIF4B and Ded1 have key but separate roles in driving the stress response Heat shock invokes rapid RNA degradation by Xrn1, selective for translation machinery

## INTRODUCTION

All organisms are subject to a continuously changing environment, to which they must adapt in order to survive. This problem is especially acute for unicellular, non-motile organisms such as the budding yeast *Saccharomyces cerevisiae*. In general, budding yeast respond to stress by inducing global changes in gene expression. At the transcriptional level, this involves the activation of the environmental stress response, in which hundreds of stress-response genes are upregulated, and genes encoding ribosome maturation and protein synthesis factors are suppressed. To a large extent, these coordinated changes in gene expression are induced regardless of the identity of the initiating stress (Gasch et al., 2000).

Transcriptional reprogramming is complemented with rapid posttranscriptional changes, particularly at the level of protein synthesis. Cytoplasmic translation is dramatically attenuated in response to a variety of environmental stresses, including various types of nutrient deprivation, but also physical stresses involving changes in temperature, osmotic balance, or oxidation state. In terms of both speed and scale, glucose starvation triggers the most drastic translational shutdown of any stress (Ashe et al., 2000; Kuhn et al., 2001). Glucose is the preferred energy and carbon source for yeast, and its absence quickly reduces cellular biosynthetic capacity. Physical stresses such as heat shock can similarly limit biosynthesis, while also damaging the existing proteome. In response, cells halt bulk protein synthesis until protective measures are in place.

Translation initiation is generally the rate-limiting step in protein synthesis, and a frequent target of regulation, including during stress; reviewed in (Crawford and Pavitt, 2019; Dever et al., 2016; Janapala et al., 2019). The initiation process begins with assembly of the 43S pre-initiation complex (PIC), comprised of the 40S subunit plus eIF1, eIF1A, eIF3, eIF5, and eIF2:GTP in complex with the initiator tRNA (Hinnebusch, 2017). In parallel, the mRNA is activated for translation by the eIF4F complex, consisting of the cap-binding protein eIF4E, the scaffolding subunit eIF4G, and the ATP-dependent helicases eIF4A and Ded1 (Gao et al., 2016). The PIC is recruited to the 5′ end of the mRNA with the help of the eIF4F complex, eIF3, and eIF4B (Mitchell et al., 2010; Park et al., 2011; Sen et al., 2015; Walker et al., 2013), and begins scanning along the transcript until it reaches the start codon. The scanning process is aided by the helicase activities of Ded1 and eIF4A, which help unwind secondary structure ahead of the translocating ribosome. Ded1 is preferentially required for translation of mRNAs with highly structured 5′ UTRs, whereas eIF4A is required for optimal translation of all mRNAs, regardless of secondary structure (Iserman et al., 2020; Sen et al., 2015). eIF4B may also assist in the scanning process through its stimulatory effect on eIF4A (Andreou et al., 2017; Sen et al., 2016; Sen et al., 2015; Walker et al., 2013).

The best studied example of translational control during stress involves the heterotrimeric eIF2 complex, which delivers the initiator tRNA to the 43S PIC. Amino acid starvation triggers the activation of the Gcn2 kinase, whose sole target is a conserved serine residue of eIF2α (Dever et al., 1992; Dey et al., 2005; Harding et al., 2000). Phosphorylated eIF2α blocks recycling of the complex for use in subsequent rounds of translation, and thus impairs bulk translation initiation. The Gcn2-eIF2α pathway is highly conserved throughout eukaryotes (Castilho et al., 2014) but, at least in yeast, it is activated in only a limited number of stress conditions; reviewed in (Simpson and Ashe, 2012). Notably, eIF2α phosphorylation is not required for translation inhibition in response to glucose withdrawal (Ashe et al., 2000) or heat shock (Grousl et al., 2009). The mechanism of translational arrest during these stresses therefore remained unclear.

Historically, a powerful tool for analyzing protein-RNA interactions, including those involved in translation, has been ultraviolet (UV) crosslinking (Dreyfuss et al., 1984). In vivo irradiation with UV light induces covalent crosslinks between protein and RNA. Subsequently, specific purification of a given protein allows for the identification of bound RNAs (the CRAC and CLIP methods). In a reciprocal approach, the RNA itself can be used to pull down associated proteins. Polyadenylated transcripts can be purified using oligo(dT) selection (Castello et al., 2012; Dreyfuss et al., 1984; Dreyfuss et al., 1993; Perez-Perri et al., 2018), but this approach is not applicable to the majority of eukaryotic RNAs, including immature mRNAs, rRNA, tRNA, and a host of additional noncoding RNAs. More recently, alternative methods have been developed to purify RNA regardless of class (Bao et al., 2018; Huang et al., 2018; Queiroz et al., 2019; Shchepachev et al., 2019; Trendel et al., 2019; Urdaneta et al., 2019). One such technique is TRAPP (total <u>R</u>NA-<u>a</u>ssociated <u>p</u>roteome <u>p</u>urification), which takes advantage of the intrinsic affinity between RNA and silica to isolate crosslinked protein-RNA complexes (Shchepachev et al., 2019).

Here, we applied TRAPP to yeast cells exposed to either glucose withdrawal or heat shock. This revealed extensive remodeling of the yeast protein-RNA interactome in response to stress. Translation initiation factors specifically involved in the recruitment and scanning process, eIF4A, eIF4B, and Ded1, dissociate from mRNA during either stress, suggesting that translation is repressed by a common mechanism. We used crosslinking and sequencing analysis to identify the precise RNA binding sites for each factor before and after stress exposure. These analyses showed that translation initiation is completely abrogated within the first 30 sec of glucose withdrawal, while heat shock induces a more gradual response. Finally, we show that the shutdown in translation initiation during heat shock results in mRNA degradation. This is selective for components of the translation machinery, presumably enforcing the translation shutdown. Taken together, this work provides new insights into stress responses and the mechanism of translational repression.

## RESULTS

### Global RBP dynamics in response to cell stress

We previously developed TRAPP as a method to characterize the global RNA-binding proteome (Shchepachev et al., 2019). Here, we applied TRAPP to assess RBP dynamics in response to glucose starvation or heat shock. An overview of the approach is shown in Fig. 1A. Yeast cells were grown in the presence of the photoreactive nucleobase 4-thiouracil (4tU), which is incorporated into nascent RNA during transcription. Following labeling, the cultures were rapidly filtered and shifted to medium containing the nonfermentable carbon sources of glycerol and ethanol (glucose withdrawal) or to standard glucose medium pre-warmed to 42°C (heat shock). At defined time points after transfer (2, 4, 8, 12, and 16 minutes), cells were irradiated with 350nm UV light to induce crosslinks between 4tU-labelled RNAs and interacting RBPs. For comparison, cells were also irradiated prior to transfer (control), or transferred and irradiated without being subjected to stress (mock treated).

**Figure 1.**
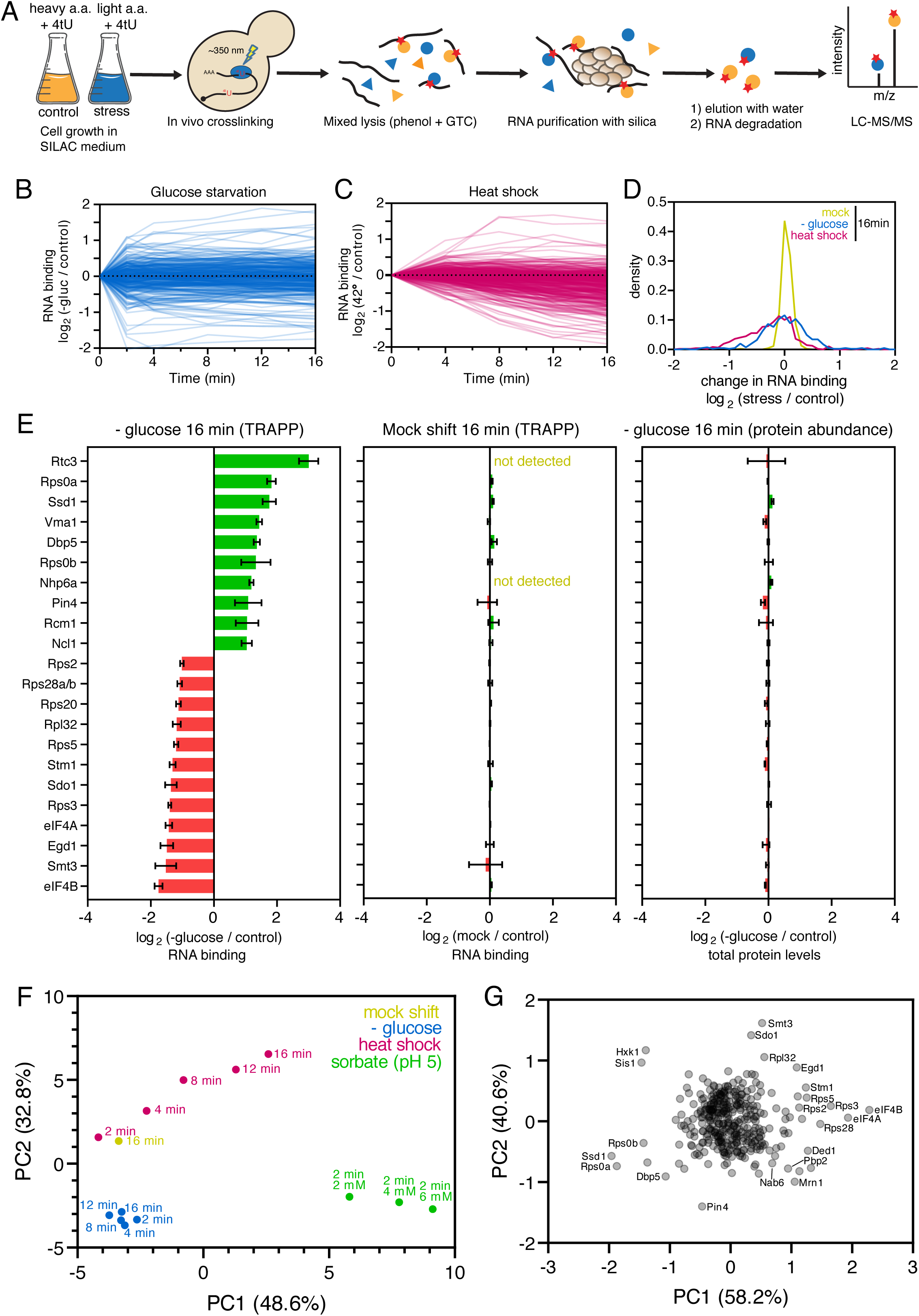
The impact of stress on the yeast RNA binding proteome. (A) Summary of the TRAPP protocol. See main text for details. (B) Time course showing changes in RNA association during glucose starvation for individual RBPs. (C) Same as (B) but for heat shock. (D) Density plot showing changes in RNA binding at 16 min following a mock shift, glucose starvation, or heat shock. (E) Bar chart showing all proteins with greater than 2-fold change in RNA association after 16 min of glucose starvation (*left)* or a mock shift *(center)*. The right-hand panel shows changes in protein abundance following 16 min of glucose starvation *(right)*. (F) Principal component analysis (PCA) showing differences between conditions and timepoints. Axis titles show the extent of variation explained by a given principal component. (G) PCA comparing the changes in RNA binding for individual proteins following heat shock, glucose starvation, or mock shift of 16 min. See also Fig. S1.

To quantify protein association with RNA, the TRAPP protocol also incorporates stable isotope labeling in cell culture (SILAC). Control cells were grown in media containing _13_C_6_ (‘heavy’) arginine and lysine, while stressed cells were cultured with standard amino acids (‘light’). Stressed and control cultures were combined in equal proportion following irradiation and lysed together under denaturing conditions. Subsequently, the cleared cell lysate was incubated with silica beads, which bind RNA along with any crosslinked protein. Following elution of the RNA:protein complexes, the RNA component was degraded, and the remaining protein was analyzed by mass spectrometry.

After filtering for proteins that were previously identified as high-confidence RNA-binders (Shchepachev et al., 2019), we quantified the association of 338 and 399 proteins across all time points following glucose starvation and heat shock, respectively (Fig. 1B-C). Most RBPs showed similar RNA-association before and after glucose withdrawal; in total, only 22 proteins showed a greater than 2-fold shift in RNA binding by 16 min (Fig. 1E). Heat shock induced a more extensive response, with a greater bias towards loss of RNA-binding (Figs. 1C-D, S1A). In contrast to the two stresses, a mock shift for 16 min produced no substantial changes in RNA binding (Fig. 1D-E, S1A). As an additional control, we also measured total protein levels before and after each stress (Figs. 1E, S1B). RBP abundance was generally constant, suggesting that the observed differences in TRAPP recovery are attributable to changes in RNA association.

In general, glucose starvation caused more rapid changes in the RNA-protein interactome compared to heat shock. Many RBPs changed dramatically within the first two minutes following glucose depletion, and were unchanged thereafter (Fig. 1B and S1D). By contrast, heat shock induced more gradual, progressive changes in RNA binding throughout the time course (Fig. 1C). These observations were further supported by principal component analysis (PCA; Fig. 1F). All of the glucose starvation time points clustered close together, indicating a high-degree of similarity following the initial rapid response. With heat shock, individual data points were more distinct in the PCA, showing a clear progression throughout the time course.

A decrease in intracellular pH has been suggested to underlie the response of yeast cells to multiple stresses (Dechant et al., 2014; Garcia et al., 2017; Munder et al., 2016). We therefore tested the effects of sorbic acid, a protonophore which equilibrates the cytosolic and extracellular pH. Cell cultures were incubated for 2 min in medium buffered at pH 5, together with increasing concentrations of sorbic acid (2, 4, and 6 mM) that are expected to correlate with decreasing intracellular pH (Munder et al., 2016). We observed substantial remodeling of the RBPome following sorbic acid treatment, but the changes were distinct from either glucose starvation or heat shock (Fig. 1F, S1C). We conclude that pH-mediated signaling is not integral to the changes in RNA binding observed during these stresses. However, we cannot exclude the possibility that sorbate treatment induces stress effects in addition to the change in pH, and these data were not extensively analyzed.

PCA analyses were also used to assess changes across the proteome for the glucose withdrawal and heat shock data (Fig. 1G). A relatively small number of proteins were clearly outliers in their response to stress. Specific RBPs will be discussed below, but we note that a group of proteins showing strongly altered RNA binding, particularly Pin4, Mrn1, Pbp2, and Nab6, remain relatively uncharacterized (Hogan et al., 2008; Riordan et al., 2011). Their identification shows the potential value of TRAPP in providing initial functional data on uncharacterized factors. Further analyses of these proteins will be reported elsewhere.

### Glucose starvation and heat shock have distinct effects on ribosome biogenesis

Given the very high rate of yeast ribosome synthesis, TRAPP datasets are highly enriched for factors involved in ribosome maturation (Shchepachev et al., 2019). Analysis of these RBPs during stress revealed several noteworthy patterns. Strikingly, most ribosome maturation factors showed no appreciable drop in RNA binding in response to glucose starvation (Fig. 2A). This is consistent with prior observations that glucose withdrawal rapidly pauses ribosome maturation and stabilizes the pre-rRNA (Abramczyk and Tollervey, unpublished observations). A substantial drop in RNA binding was seen only for Sdo1, which catalyzes the final step of 60S maturation (Kargas et al., 2019) (Fig. S1D). Heat shock, by contrast, induced a progressive decrease in RNA binding over the 16 min time course for nearly all maturation factors. This indicates that the inhibition of ribosome synthesis develops over time, with maturation or degradation of the nascent particles. Notably, Sdo1 was again one of the few exceptions, showing a modest, but significant, increase in RNA association. This may indicate some accumulation of very late pre-ribosomes immediately prior to final maturation of 60S subunits.

**Figure 2.**
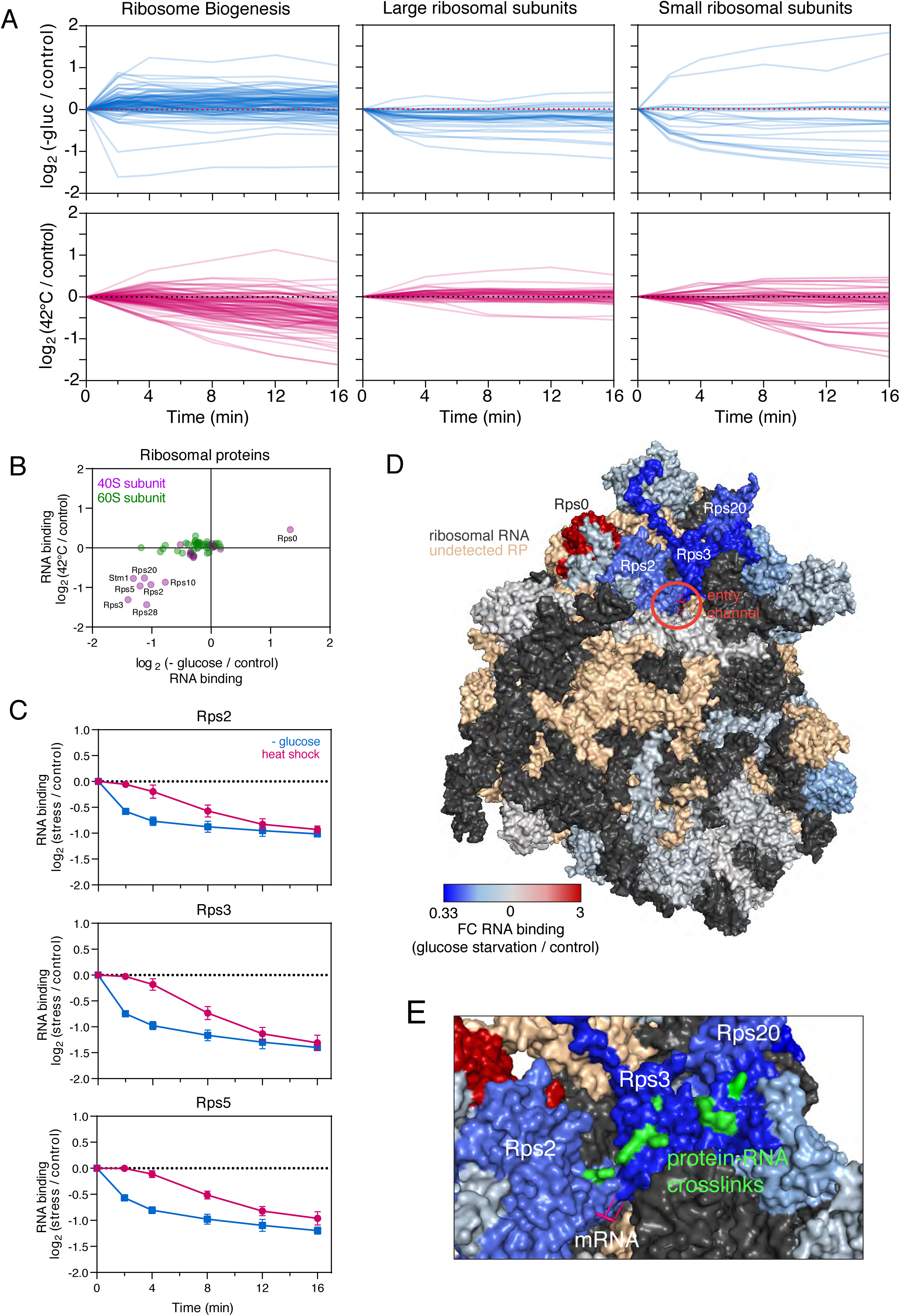
Changes in RNA binding among ribosomal proteins. (A) Time course showing changes in RNA binding during glucose withdrawal *(upper)* and heat shock *(lower)* for various classes of RBPs. Included in the figure are ribosome biogenesis factors *(left)*, large ribosomal subunits *(center)*, and small ribosomal subunits *(right)*. (B) Scatterplot comparing the effects of glucose starvation and heat shock at 16 min on RNA binding for ribosomal proteins. (C) Time course showing RNA association for Rps2, Rps3, and Rps5. (D) Crystal structure (3J77) of the yeast ribosome highlighting the changes in RNA association for each detected ribosomal protein. (E) A closeup view of the mRNA entry channel with amino acid-RNA crosslinking sites highlighted in green.

### Ribosomal protein binding dynamics in response to cell stress

We next turned our attention to ribosomal proteins (RPs), a number of which showed decreased RNA association following either glucose withdrawal or heat shock (Fig. 2A). Direct comparison of the two datasets revealed that a similar collection of RPs from the 40S subunit were altered in each stress (Fig. 2B). However, in each case, glucose starvation induced a more rapid response (for examples see Fig. 2C). We mapped these TRAPP results onto the structure of the ribosome, with each protein colored according to its change in RNA binding at 16 min following glucose withdrawal (Fig. 2D). Intriguingly, RPs with altered binding predominately clustered around the mRNA channel. Indeed, the subset of proteins which directly contact the translating mRNA (e.g. Rps3) showed the most substantial drop in RNA association. We conclude that decreased translation during stress drives reduced RNA association for RPs that would otherwise contact the mRNA.

Finally, we mapped precise amino acid sites of RNA crosslinking to RPs, using published data from the iTRAPP method (Shchepachev et al., 2019). Visualization of these sites on the ribosomal structure (Fig. 2E), revealed a succession of crosslinked amino acids along the surfaces of Rps2 and Rps3, apparently tracing the path of the mRNA as it approaches the channel. To our knowledge, RNA binding in this region has not been observed previously using crystallography or cryoEM, presumably due to structural flexibility in the mRNA. This highlights the utility of TRAPP for capturing flexible or transient interactions.

### A common set of translation initiation factors are regulated in response to stress

We next asked whether the TRAPP data could shed light on the mechanism of translational repression. We focused first on translation initiation, as it is the most common target of regulation. The RNA-binding dynamics for each eukaryotic initiation factor (eIF) are shown in Fig. S3. An overview of the translation initiation pathway is shown in Fig. 3A (for details, see Introduction). Each protein is colored according to its change in RNA binding, specifically in response to glucose withdrawal, but most eIFs showed a consistent response to both stress conditions (Fig. S3).

**Figure 3.**
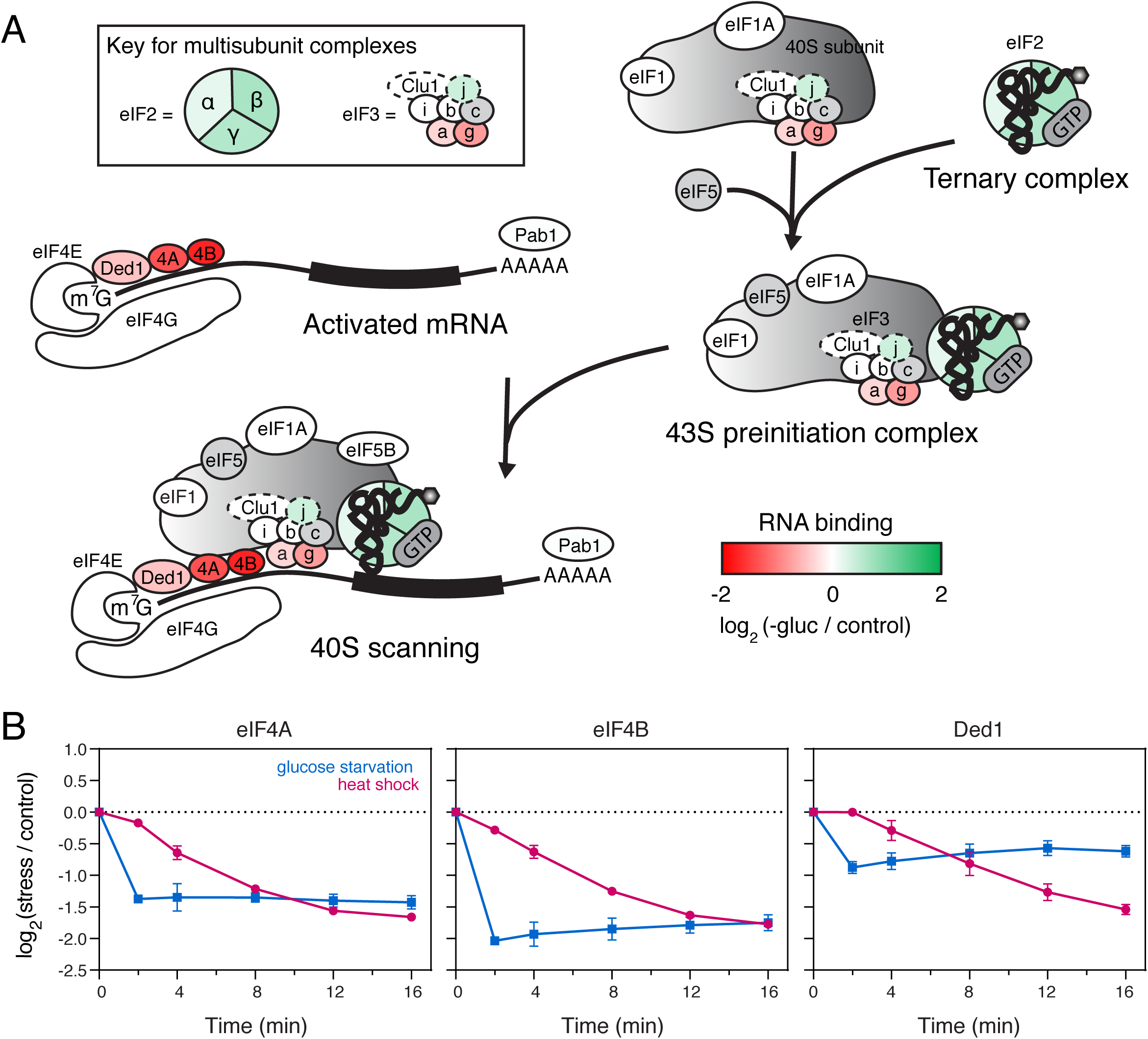
Changes in RNA binding among translation initiation factors. (A) Overview of the translation initiation process. Each protein is colored according to its change in RNA association during glucose starvation. Translation initiation factors shown in grey were not detected as RNA-binding. (B) Time course showing changes in RNA binding for eIF4A, eIF4B, and Ded1 following glucose starvation (blue) and heat shock (pink). See also Fig. S2.

We first considered the set of initiation factors primarily associated with the 40S subunit, including eIF1, eIF1A, eIF5, and eIF5B. For this group, each protein remained associated with RNA following stress (Fig. 3A, S3A), suggesting that their binding to the 40S subunit was unaltered. Next, we considered the eIF3 complex, subunits of which predominately bind either the mRNA or 40S ribosomes. The eIF3g and eIF3a subunits both contact the mRNA during translation (Valasek et al., 2017) and showed decreased RNA-association. Conversely, eIF3b and eIF3i, which only interact with the 40S ribosome, were unchanged. The enigmatic Clu1 subunit was also unchanged in RNA binding, but its place in the complex, if any, is unclear (Vornlocher et al., 1999). Only eiF3j showed increased RNA association following both stresses. However, eIF3j is not a constitutive subunit of the eIF3 complex and, despite its name, has only been implicated in translation termination (Young and Guydosh, 2019).

These findings suggested that the 43S pre-initiation complex assembles normally during stress. However, a downstream block in translation initiation may impair its recruitment to mRNA. We therefore assessed mRNA-specific initiation factors that recruit the 43S complex. The cap binding protein eIF4E, was largely unaffected by either stress (Fig. S3A), whereas RNA binding by eIF4G was unaltered by glucose withdrawal, but modestly reduced with heat shock. More dramatic effects on RNA binding were seen for the scanning factors eIF4A and eIF4B (Fig. 3B). RNA association was strongly decreased following either stress, but with much faster kinetics during glucose withdrawal. These observations are consistent with the reported loss of eIF4A from polysomes following glucose withdrawal (Castelli et al., 2011). The eIF4A-homolog Ded1 also showed a robust decrease in RNA binding during both stresses, though this was significantly less pronounced for glucose withdrawal (discussed in more detail below).

When bound to mRNA, eIF4A, eIF4B, and Ded1 recruit the 43S preinitiation complex, and Ded1 additionally assists in scanning and start codon recognition (Andreou et al., 2017; Guenther et al., 2018; Sen et al., 2015; Walker et al., 2013). Loss of RNA binding by these factors is therefore expected to strongly impair translation initiation. We conclude that a specific block in PIC recruitment and/or mRNA scanning underlies the translation repression seen following either glucose withdrawal or heat shock.

### Differential RNA binding by eIF4A, eIF4B, and Ded1 upon stress

To further investigate the role of eIF4A, eIF4B, and Ded1 in translation shutoff, we mapped the RNA binding sites for each protein using crosslinking and analysis of cDNA (CRAC). Strains were constructed in which eIF4A and Ded1 were expressed as N-terminal, FH-tagged (Flag-Ala_4_-His_8_) fusion proteins and eIF4B was expressed with a C-terminal HF tag (His_8_-Ala_4_-Flag), under control of the endogenous promoters. The fusion proteins each supported wild-type growth, indicating that they are functional (data not shown). Actively growing cells expressing the fusion proteins were UV-irradiated at 254 nm for ∼4-6 sec in a VariX crosslinker to covalently fix direct protein:RNA contacts. After stringent, tandem-affinity purification, partial RNase digestion, and radiolabeling, protein:RNA complexes were isolated using SDS-PAGE (Fig. S4A). Subsequently, crosslinked RNA fragments were amplified using RT-PCR and analyzed by high-throughput sequencing. For each protein, we collected datasets from unstressed cells (‘control’), mock-shifted cells without stress (16 min), and following glucose withdrawal (30 sec and 16 min) or heat shock (16 min) (Fig. S4B). Metaplots of individual replicates showed good reproducibility (Fig. S4D) and for subsequent analyses, replicate datasets were merged to provide improved coverage along individual transcripts.

We first examined interactions between each translation factor and ribosomal RNA in control cells (Fig. S5A-B). eIF4A showed weak binding throughout 18S, together with two sharp peaks in 25S. However, both crosslinking sites were buried within the ribosome, so likely represent sequencing artifacts. Clear results were seen with eIF4B, for which we observed a single major crosslinking site, situated close to the mRNA exit channel (nucleotide 1,060). Ded1 also crosslinked mainly to 18S rRNA, with prominent peaks near the mRNA entry (nucleotide 492) and exit (nucleotide 1053) channel, and additional binding at position 719. Notably, crosslinking at all three sites is consistent with previously reported interactions between Ded1 and 18S rRNA (Guenther et al., 2018).

A breakdown of crosslinked RNAs by biotype revealed substantial differences pre- and post-stress. In unstressed cells, eIF4B primarily targeted mRNAs, but this enrichment was abruptly lost following exposure to either stress (Fig. 4A). By contrast, cells subjected to a mock shift were indistinguishable from the control, indicating that the experimental protocol per se did not significantly perturb the cells. Similar, but less dramatic, trends were seen for Ded1 and eIF4A (Fig. S4C). Analysis of binding sites on individual mRNAs provided a high-resolution snapshot of translation dynamics (Fig. 4B-D). The vast majority of transcripts showed decreased binding by eIF4B, Ded1, and eIF4A upon exposure to stress. Remarkably, binding was greatly decreased within the first 30 sec of glucose withdrawal, consistent with prior reports that translation initiation is repressed within the first minute (Ashe et al., 2000). The reduction was even more pronounced when we considered only 5′ binding, defined as reads mapping to either the 5′ UTR or the first 150 nucleotides of the open reading frame (Figs. 4B-D). This specific loss of 5’ binding was also seen in a metagene analysis of the top 2,000 bound mRNAs (Figs. 4E and S4D). Heatmaps of the distribution of eIF4B along each of the 2,000 mRNAs confirmed that loss of 5’ binding is a general feature (Fig. S4E).

**Figure 4.**
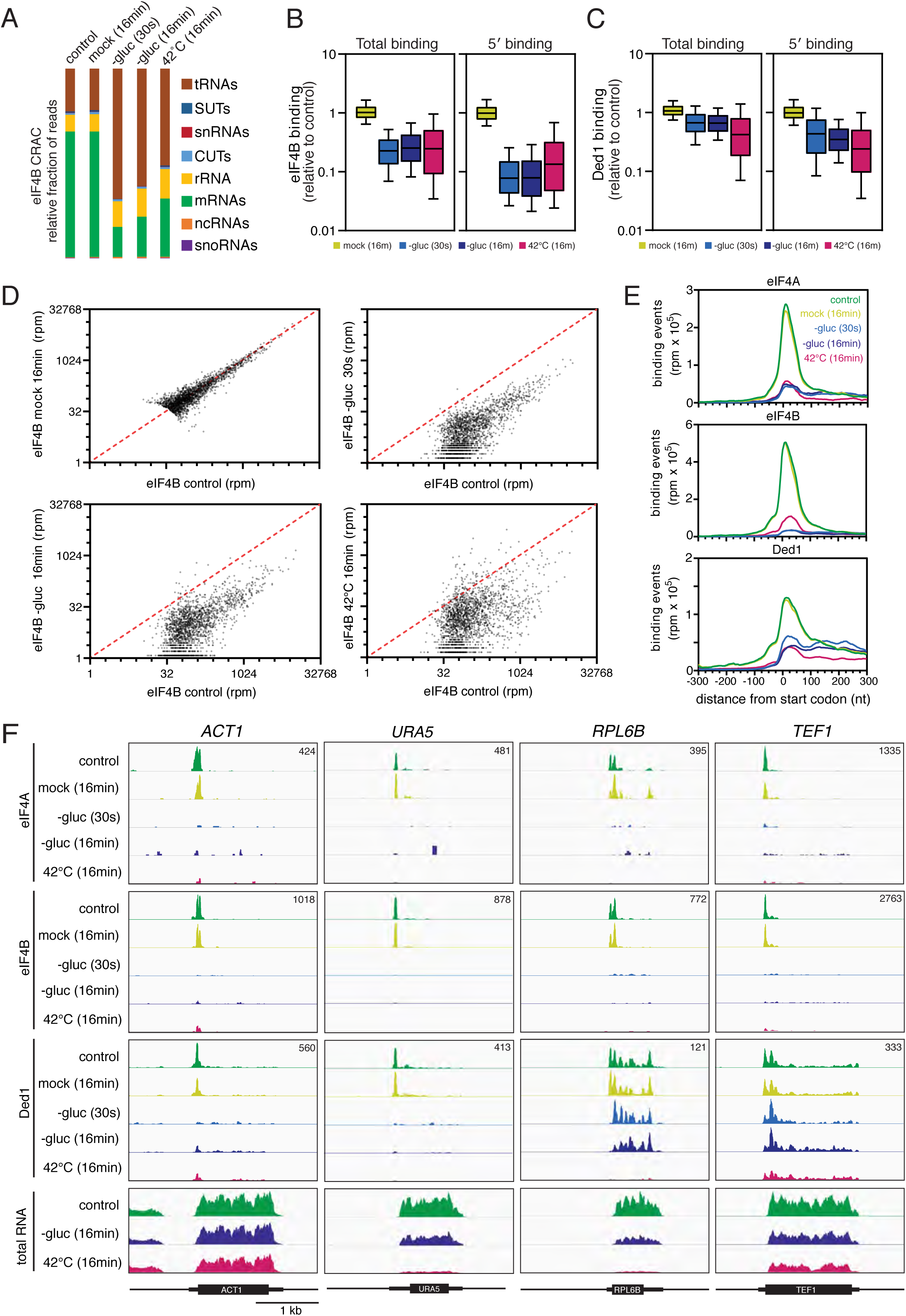
Genome-wide analysis of the RNA binding profiles of eIF4A, eIF4B, and Ded1. (A) Breakdown of eIF4B-bound RNAs by biotype. (B) Boxplot showing the changes in eIF4B binding to individual mRNAs following either mock shift or stress. Each box represents the median with 25^th^ and 75^th^ percentiles. The whiskers show the 10^th^ and 90^th^ percentiles. For panels (B-E), all analyses are based on a set of 2,000 transcripts which show the strongest binding to eIF4B in unstressed conditions. (C) Same as (B) but for Ded1. (D) Scatter plots comparing the changes in RNA binding between control and either mock shift (16 min), glucose starvation (30 sec), glucose starvation (16 min), or heat shock (16 min). (E) Metaplots showing the distribution of eIF4A, eIF4B, and Ded1 binding around the mRNA start codon. (F) Binding of eIF4A, eIF4B, and Ded1 across the *ACT1, URA5, RPL6B*, and *TEF1* mRNAs. Each set of tracks is normalized to total library size using reads per million, with the exact value indicated in the upper right corner of each box. RNAseq traces are shown at the bottom as a control. Each track is normalized to a spike-in control, and thus represents the absolute abundance of each mRNA compared to the control. Each box represents a 3 kb window; a scale bar is shown at the bottom. See also Figs. S3 – S5.

Having examined the ‘average’ binding profile by metaplot, we next investigated how binding varied between individual mRNAs (Figs. 4F and S5A). On most mRNAs, eIF4A and eIF4B targeted a single site close to the start codon (shown for *TEF1* and *URA5*). For a minority of transcripts, we observed two distinct peaks, usually on either side of the start codon (shown for *ACT1* and *RPL6B*). Overall, the two translation factors displayed a strikingly similar binding profile, consistent with the role of eIF4B as a cofactor for eIF4A (Andreou et al., 2017). In agreement with the metagene analysis, most individual transcripts showed sharply reduced binding to both eIF4A and eIF4B following stress.

Ded1 displayed a more complicated binding pattern. For most transcripts, including *ACT1* and *URA5* (Fig. 4F), Ded1 showed a similar distribution to eIF4A and eIF4B, binding at a single site near the 5′ end in unstressed cells, and largely dissociating during stress. However, a substantial fraction of transcripts showed pervasive binding throughout the length of the mRNA. To quantify this difference, we generated heatmaps in which individual transcripts were sorted by their ratio of 5′ versus pervasive binding. For Ded1, 66% of transcripts showed at least as much binding at downstream sites as at the 5′ end (Fig. S5B, S5D). By contrast, only 5% of transcripts showed a comparable ratio for eIF4B (Fig. S5C-D). The *RPL6B* mRNA illustrates these points well (Fig. 4F). In unstressed cells, Ded1 was bound throughout the transcript, with modest enrichment at the 5′ end. Intriguingly, 5′ binding was selectively lost in response to glucose withdrawal, while downstream binding was maintained. By contrast, heat shock resulted in a general loss of Ded1 binding across the length of the mRNA. Similar results were seen for other transcripts, including *TEF1* (Fig. 4F) and *RPL34B* (Fig. S5A), albeit to varying degrees. At the metagene level, we observed a specific reduction in 5′ binding, while downstream binding was relatively unaltered following glucose withdrawal but decreased following heat shock (Fig. 4E). Importantly, these observations are consistent with the TRAPP data, which showed that glucose withdrawal had a relatively modest effect on binding of Ded1 to RNA in comparison to heat shock (Fig. 3B).

For most mRNA species, targeting by Ded1 was approximately proportional to transcript abundance, but there were a number of outliers (Fig. S5E). The most prominent was the *DED1* mRNA itself, which was highly bound by Ded1. Intriguingly, most binding was concentrated within the 3′ UTR (Fig. S5A), a pattern counter to most other mRNAs, and suggestive of some form of auto-regulatory control.

### Heat shock triggers the degradation of translation-associated mRNAs

Our results indicate that specific translation initiation factors dissociate from mRNAs in response to stress. To control for changes in mRNA levels, we harvested RNA before and 16 min after each stress and performed RNAseq, with RNA from *Schizosaccharomyces pombe* included as a spike-in control for quantitation. Two replicates were collected for each condition and these showed excellent reproducibility (Fig. S6). Glucose withdrawal had inconsistent, but relatively mild effects on overall mRNA abundance, whereas heat shock induced a modest global decrease (down ∼25%) in mRNA levels (Fig. 5A). Notably, the reduction in total mRNA was much less than the reduction in RNA binding by eIF4A, eIF4B, and Ded1 (Fig. 3B), showing that the decreased initiation factor binding was not due to reduced mRNA abundance.

**Figure 5.**
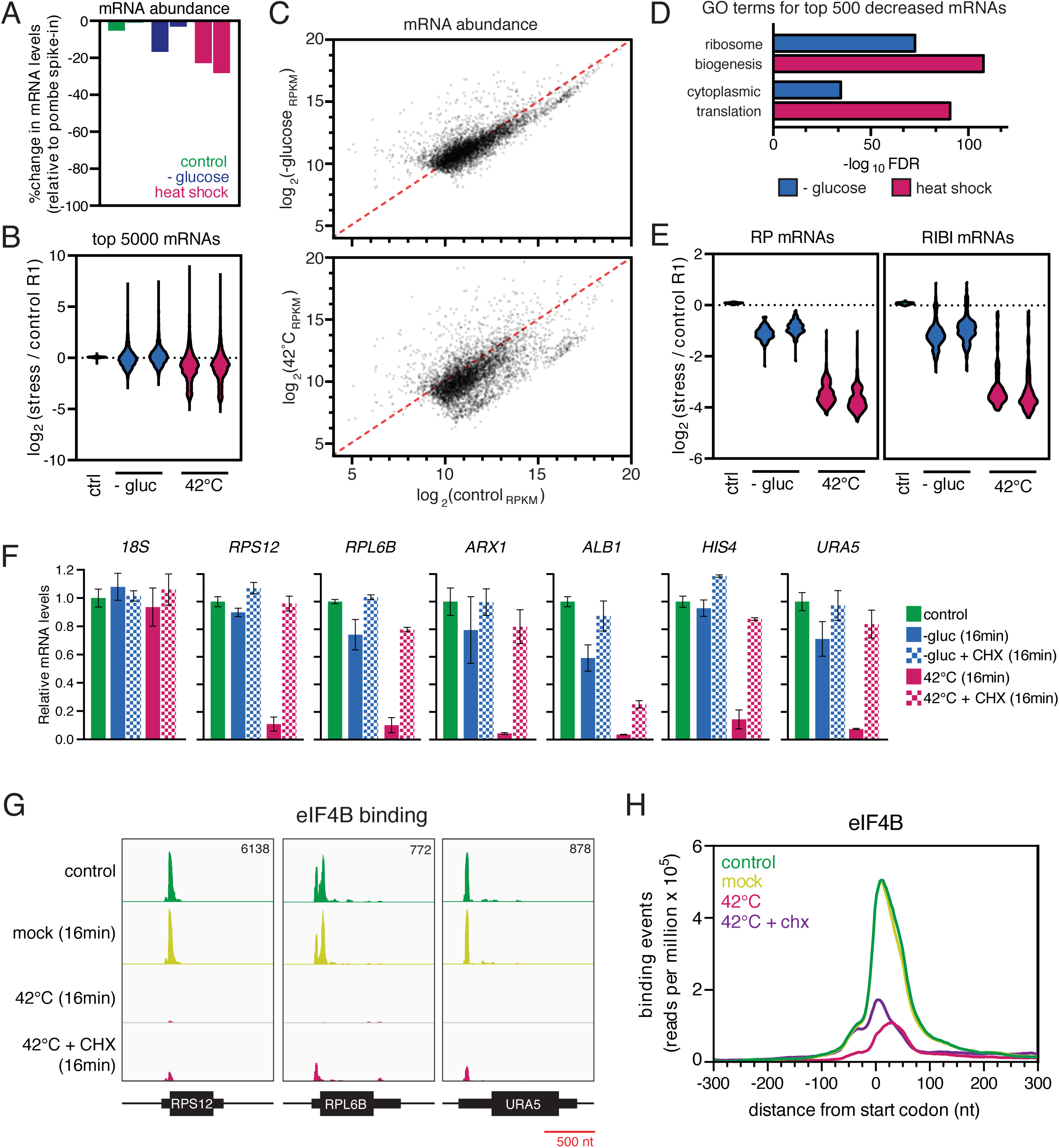
Global analysis of mRNA levels in response to stress. (A) Bar graph showing the change in total mRNA abundance relative to an *S. pombe* spike-in control. (B) Violin plots of the 5,000 most-abundant mRNAs showing changes in transcript levels in response to stress. Expression levels are calculated relative to control replicate #1. (C) Scatter plots comparing mRNA levels in glucose starvation *(upper)* or heat shock *(lower)* relative to control. Each plot includes the 5,000 most-abundant mRNAs. (D) GO term enrichment among the 500 most decreased mRNAs for each stress. (E) Violin plots showing the changes in mRNA levels for ribosomal protein (RP) mRNAs *(left)*, or ribosome biogenesis (RIBI) mRNAs *(right)*. (F) Bar graphs showing changes in mRNA levels following stress with and without co-treatment with the translation elongation inhibitor cycloheximide (CHX). Each bar represents the average of two independent experiments, and the error bars show the standard deviation. (G) Binding of eIF4B across selected transcripts in control cells, following a mock shift, heat shock, or heat shock in the presence of cycloheximide. (H) Metaplots showing the distribution of eIF4B binding around the mRNA start codon. See also Fig. S6.

This conclusion was supported by analysis of individual mRNAs (Fig. 5B-C). Most mRNAs were only mildly affected by glucose depletion, but a substantial fraction of transcripts were sharply decreased in response to heat shock. GO analysis of the 500 most-depleted mRNAs revealed strong enrichment for translation-associated transcripts, primarily components of the ribosome biogenesis and cytoplasmic translation machinery (Fig. 5D). This enrichment was seen for both stresses, but was more significant following heat shock. We confirmed these results with quantitation of mRNAs that either encode ribosomal proteins or fall into the <u>ri</u>bosome <u>bi</u>ogenesis (RIBI) regulon (Fig. 5E), a group of over 200 coordinately regulated factors (Jorgensen et al., 2004; Klinge and Woolford, 2019; Wade et al., 2006). Both groups of mRNAs were, on average, reduced <2 fold by glucose depletion and ∼8 fold following heat shock (Fig. 5E).

The stability of a given mRNA species is often determined by a competition between translation and mRNA degradation (Chan et al., 2018; Huch and Nissan, 2014; Schwartz and Parker, 1999). We therefore tested whether the decrease in abundance of specific transcripts could reflect mRNA decay induced by the translation shutdown. We analyzed the abundance of six transcripts by RT-qPCR and compared them to 18S rRNA (Fig. 5E). Consistent with the RNAseq data, glucose withdrawal was associated with only modest changes (<2 fold), whereas heat shock was accompanied by ≥10 fold reductions in mRNA abundance. To determine the role of translation, heat shock was combined with cycloheximide treatment, which inhibits translation elongation, freezing ribosomes in place. Notably, treatment with cycloheximide largely prevented the drop in mRNA levels for most transcripts (Fig. 5F). We conclude that heat shock-induced mRNA decay requires continued translation elongation. In the absence of cycloheximide, mRNA degradation may be triggered by the shutdown in translation initiation combined with the ensuing ribosome runoff, resulting in “naked” mRNAs susceptible to degradation factors.

One prediction of this hypothesis is that translation initiation factors dissociate from mRNAs prior to degradation of the transcript. To test this idea, we examined eIF4B binding after heat shock, either with or without cotreatment with cycloheximide. If translation factors dissociate prior to mRNA decay, then eIF4B should lose RNA binding during heat shock even if mRNA levels are stabilized. Indeed, eIF4B binding to *RPS12, RPL6B*, and *URA5* was sharply reduced following heat shock (Fig. 5G), even when the cells were cotreated with cycloheximide to block mRNA degradation. Similar results were observed genome-wide (Fig. 5H). We conclude that translation initiation shutoff occurs upstream of mRNA decay.

### Heat shock induces selective mRNA 5′ degradation

Finally, we investigated the factors required for heat shock-induced mRNA decay. In the cytoplasm, mRNAs can be 5′-degraded by the 5′ => 3′ exonuclease Xrn1, and/or 3′ degraded by the 3′ => 5′ exonuclease activity of the exosome, which also requires the Ski2 helicase as a cofactor. Strains lacking either Xrn1 or Ski2 are viable, although loss of both activities induces synthetic lethality (Brown et al., 2000). We separately deleted the genes encoding Xrn1 and Ski2 and tested whether the effects of heat shock were rescued in the mutant strains (Fig. 6A). For all mRNAs tested, levels were substantially restored in the *xrn1Δ* strain, but not in *ski2Δ.* We conclude that following heat-shock many mRNAs are subject to 5′ degradation, largely dependent on the activity of Xrn1.

**Figure 6.**
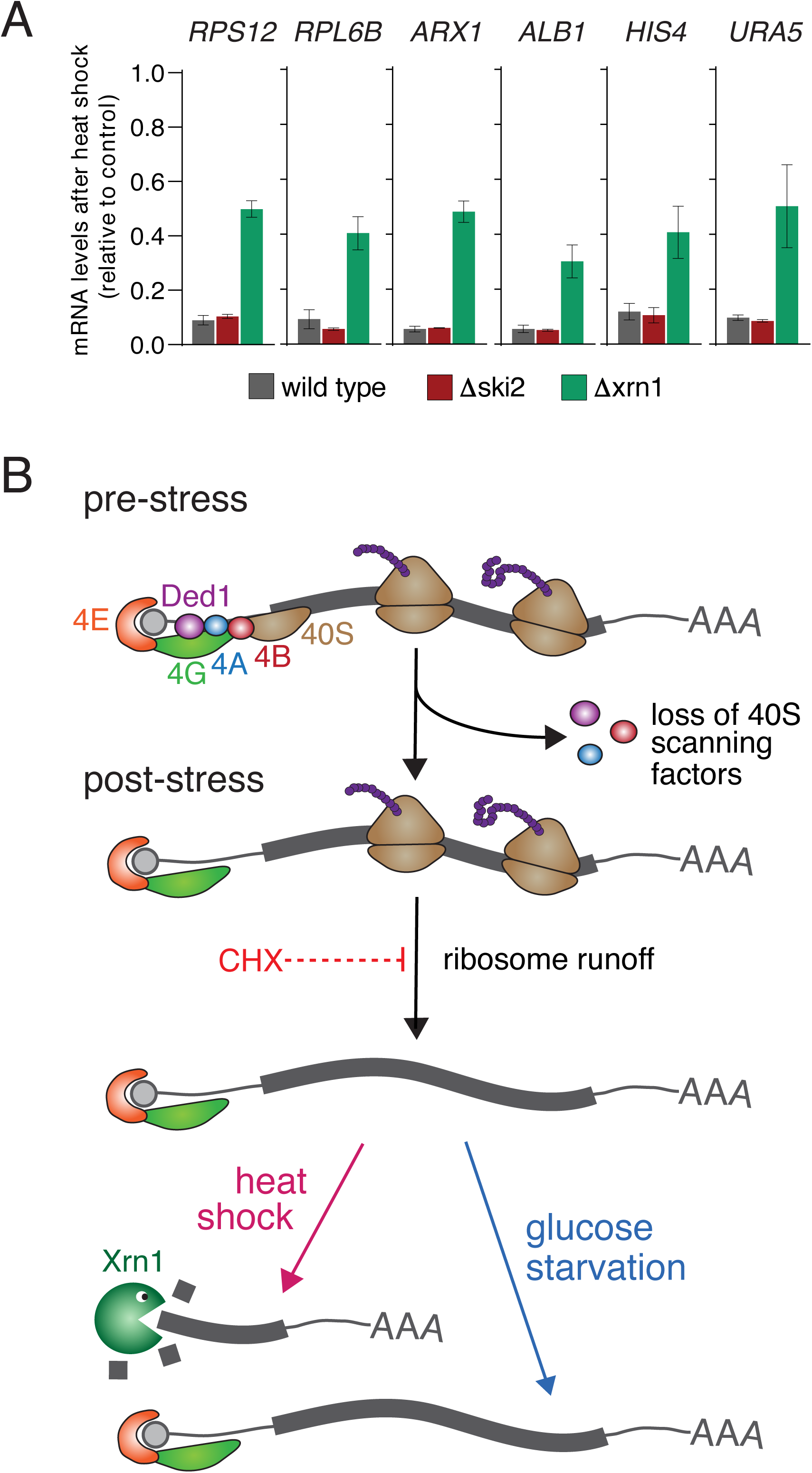
(A) Bar graph showing the changes in mRNA abundance measured by RT-qPCR following 16 min of heat shock in either wild type cells or cells deleted for *SKI2* or *XRN1*. (B) Model of the translational response to glucose starvation and heat shock. Upon exposure to either stress, the 40S scanning factors eIF4A, eIF4B, and Ded1 dissociate from the 5′-end of mRNAs, halting translation initiation. Already-initiated ribosomes continue translating before eventually terminating, leaving ‘naked’ mRNAs unprotected by the translational machinery. In the case of heat shock, Xrn1 is involved in degradation of a subset of these transcripts. With glucose starvation, by contrast, most mRNAs remain stable.

## DISCUSSION

RNA-binding proteins are critically important in the cellular response to environmental stress. The TRAPP technique allows RBP dynamics to be followed quantitatively on short time scales, and is thus well-suited for monitoring global changes in the protein-RNA interactome during stress. Here, we used TRAPP to follow RBP dynamics in *S. cerevisiae* during glucose starvation and heat shock. We observed rapid and specific changes in RNA binding for dozens of proteins, with translation-associated factors among the most significantly altered. Taken together, our results shed new light on the mechanism of translational repression in yeast.

Consistent with a general shutdown of translation in response to stress, ribosomal proteins surrounding the mRNA channel showed decreased association with RNA (Fig. 2). Similar findings have recently been reported for human cells exposed to arsenite stress (Trendel et al., 2019). In keeping with our observations, the channel proteins hRPS3 and hRPS28 showed especially strong loss of RNA binding (Trendel et al., 2019). Arsenite stress also triggered extensive ribosomal degradation, perhaps to further enforce the translation shutdown, or as a means to remove damaged ribosomes. However, we found no evidence of ribosome degradation in response to stress in yeast. Indeed, the abundance of most proteins was completely unaffected by either stress (Fig. S1), suggesting changes in RBP levels are not a significant driver of rapid translation shutoff in yeast.

Throughout eukaryotes, phosphorylation of the translation initiation factor eIF2α leads to translation shutdown in response to various stresses. However, both glucose starvation and heat shock trigger translational repression via different, but largely undefined, pathways (Ashe et al., 2000; Grousl et al., 2009). In an attempt to characterize these pathways in more detail, we examined the RNA binding dynamics of each translation initiation factor. Initiation factors specifically associated with the 40S subunit showed little change in RNA binding during either stress. Conversely, mRNA-associated factors involved in 43S complex recruitment and scanning (the RNA helicases Ded1 and eIF4A, as well as the eIF4A cofactor eIF4B) showed substantial loss of RNA binding in response to either stress. The striking similarity between the two stress conditions suggests a common mechanism of translational repression. We propose that loss of RNA binding by eIF4A, eIF4B, and Ded1 prevents recruitment and/or scanning of the 43S PIC, blocking translation initiation (Fig. 6B). Already-initiated ribosomes probably continue elongating before eventually “running off”, halting translation entirely (Ashe et al., 2000). The TRAPP data provide additional support for this ‘initiation-first’ model of translational repression. Upon glucose removal, the initiation factors eIF4A, eIF4B, and Ded1 showed an immediate decrease in RNA binding, whereas ribosomal proteins which interact with the mRNA during elongation (e.g. Rps2, 3, and 5) lost RNA binding more slowly (Fig. 2).

To understand the basis of the translation shutdown in more detail, we mapped the RNA-binding sites for eIF4A, eIF4B, and Ded1 using CRAC. In unstressed cells, eIF4A and B primarily targeted the 5′ ends of mRNAs, consistent with their role in translation initiation. Notably, binding peaked ∼20 nt downstream of the start codon, rather than within the 5′ UTR, as might be expected for scanning factors. One possible explanation is that scanning across the 5′ UTR happens relatively fast compared to the amount of time the 48S PIC spends positioned over the start codon awaiting subunit joining and subsequent translation. This is supported by ribosome profiling data which consistently shows accumulation of ribosomes at the AUG (Mohammad et al., 2019). The binding sites mapped for both eIF4B and Ded1 on the 18S ribosomal RNA place them near the exit channel (Fig. S6), suggesting each protein interacts with mRNA as it is extruded from the channel. A prolonged pause over the start codon would thus result in enhanced crosslinking downstream. We cannot, however, exclude the possibility that RNA fragments immediately adjacent to the mRNA cap may be recovered less efficiently. The short 5′ UTRs typical of budding yeast (Tuller et al., 2009), could then shift the apparent binding site downstream of the start codon.

Regardless of the precise binding site, 5′ association was clearly lost in response to stress (Fig. 4). For glucose starvation, this occurred remarkably quickly. Binding was completely ablated within just 30 sec, the earliest timepoint we were able to test. An unresolved question is how information on the depletion of glucose from the medium is gathered and transmitted to drive such rapid changes in protein binding. The proteins involved are extremely abundant; eIF4A (Tif1 plus Tif2) is present at ∼150,000 copies per yeast cell, comparable to the ribosome, with eIF4B (Tif3) and Ded1 at ∼25,000 copies (Ho et al., 2018). Presumably, extensive signal amplification is required to effectively regulate such abundant target proteins. Various signaling proteins and mRNA decay factors have been implicated in translation shutoff (Ashe et al., 2000; Coller and Parker, 2005; Holmes et al., 2004; Vaidyanathan et al., 2014), but their connections with the translation initiation machinery remain unknown.

Recent work suggests Ded1 is itself a stress-sensor due to its ability to reversibly condense into phase-separated granules in vitro and in vivo (Hondele et al., 2019; Iserman et al., 2020; Wallace et al., 2015). This accumulation is stimulated by heat shock or a drop in intracellular pH, suggesting a possible mechanism for translation inhibition (Iserman et al., 2020). The CRAC data on Ded1 association are consistent with this model, showing a general loss of interactions with mRNA translation initiation regions, but retention of binding further 3’ along the transcript (Fig. 4 and S6). The specific RNA targets for Ded1-mediated assembly into stress-induced granules remain unknown (Hilliker et al., 2011; Iserman et al., 2020), and the transcripts showing pervasive Ded1 binding are strong candidates.

Previous analyses have shown that loss of eIF4A or eIF4B in yeast confers greater inhibition of general translation than inactivation of Ded1 (Sen et 2016). Like Ded1, eIF4B also condenses into phase-separated granules during heat shock (Wallace et al., 2015). We therefore propose that they function as independent stress sensors. In principal, the loss of eIF4A binding during stress could reflect the loss of either of its reported binding partners (eIF4B or Ded1). However, the CRAC data are more consistent with eIF4B playing the major role in eIF4A recruitment.

Since heat shock and glucose withdrawal can each drive relocation of translation factors and mRNAs into phase-separated cytoplasmic granules, similar effects on RNA stability might have been anticipated. However, this is not the case. Following glucose withdrawal, mRNA levels were very stable, whereas heat shock induced a substantial decrease in abundance for many mRNAs. Depletion was particularly strong for mRNAs encoding nucleolar ribosome synthesis factors or cytoplasmic ribosomal proteins. These mRNAs were reduced 10-fold or more in 16 min, demonstrating activated RNA degradation (Fig. 5). The same mRNAs are transcriptionally repressed as part of the integrated stress response (Gasch et al., 2000), which presumably maintains prolonged repression of ribosome synthesis. Remarkably, most mRNAs were stabilized by co-treatment with the translation elongation inhibitor cycloheximide, indicating that continued elongation is required for mRNA degradation.

Taken together our findings lead us to the following model (Fig. 6B). Under normal circumstances (i.e. in the absence of cycloheximide), heat shock triggers a halt in cap-dependent translation initiation, followed by runoff of already-initiated ribosomes. Without the protection of polysomes, the resulting transcripts are subject to 5′ to 3′ degradation by the exonuclease Xrn1. By contrast, in the presence of cycloheximide, ribosomes remain associated with the transcript, and either block degradation directly, or prevent the mRNA from relocalizing to sites of decay.

This model is consistent with prior observations suggesting a competition between translation and mRNA decay. Mutations in translation initiation factors lead to increased deadenylation and decapping (Schwartz and Parker, 1999). Moreover, chemical inhibition of translation initiation results in rapid mRNA decay (Chan et al., 2018). We suggest that a similar phenomenon occurs in response to heat shock, with some transcripts subject to increased degradation in the absence of active translation. A major open question is how translation-related mRNAs are preferentially targeted during heat shock, and why they remain stable during the shift from glucose to an alternative carbon source (Fig. 5). The translation shutoff in response to glucose withdrawal is, if anything, more severe than that seen with heat shock. Cytoplasmic mRNA deadenylation is downregulated following glucose withdrawal (Hilgers et al., 2006), making it likely that glucose signaling pathways block degradation in addition to immediately blocking initiation. Identifying the key signaling factors and their downstream targets is now a priority.

## ACKNOWLEDGEMENTS

We thank Richard Clark and Angie Fawkes from the Wellcome Trust Clinical Research Facility at Western General Hospital (Edinburgh) for sequencing services. This work was supported by Wellcome through a Principle Research Fellowship to D.T. (077248), a Senior Research Fellowship to J.R. (103139) and an instrument grant (108504). TWT was supported by the Polish Ministry of Science and Higher Education Mobility Plus program (1069/MOB/2013/0). Work in the Wellcome Centre for Cell Biology is supported by a Centre Core grant (203149).

## AUTHOR CONTRIBUTIONS

SB and DT conceived the project, analyzed the data, and wrote the manuscript. SB, CS, and VS performed experiments. TT performed principal component analysis. All authors edited and reviewed the manuscript.

## DECLARATION OF INTERESTS

The authors declare no competing interests.

## DATA AVAILABILITY

The GEO accession number for all sequence data reported in this paper is GSE148166. The proteomics data are available through the ProteomeXchange Consortium via the PRIDE (Perez-Riverol et al., 2019) partner repository with the dataset identifier PXD019141.

## FIGURE LEGENDS

**Figure S1.**
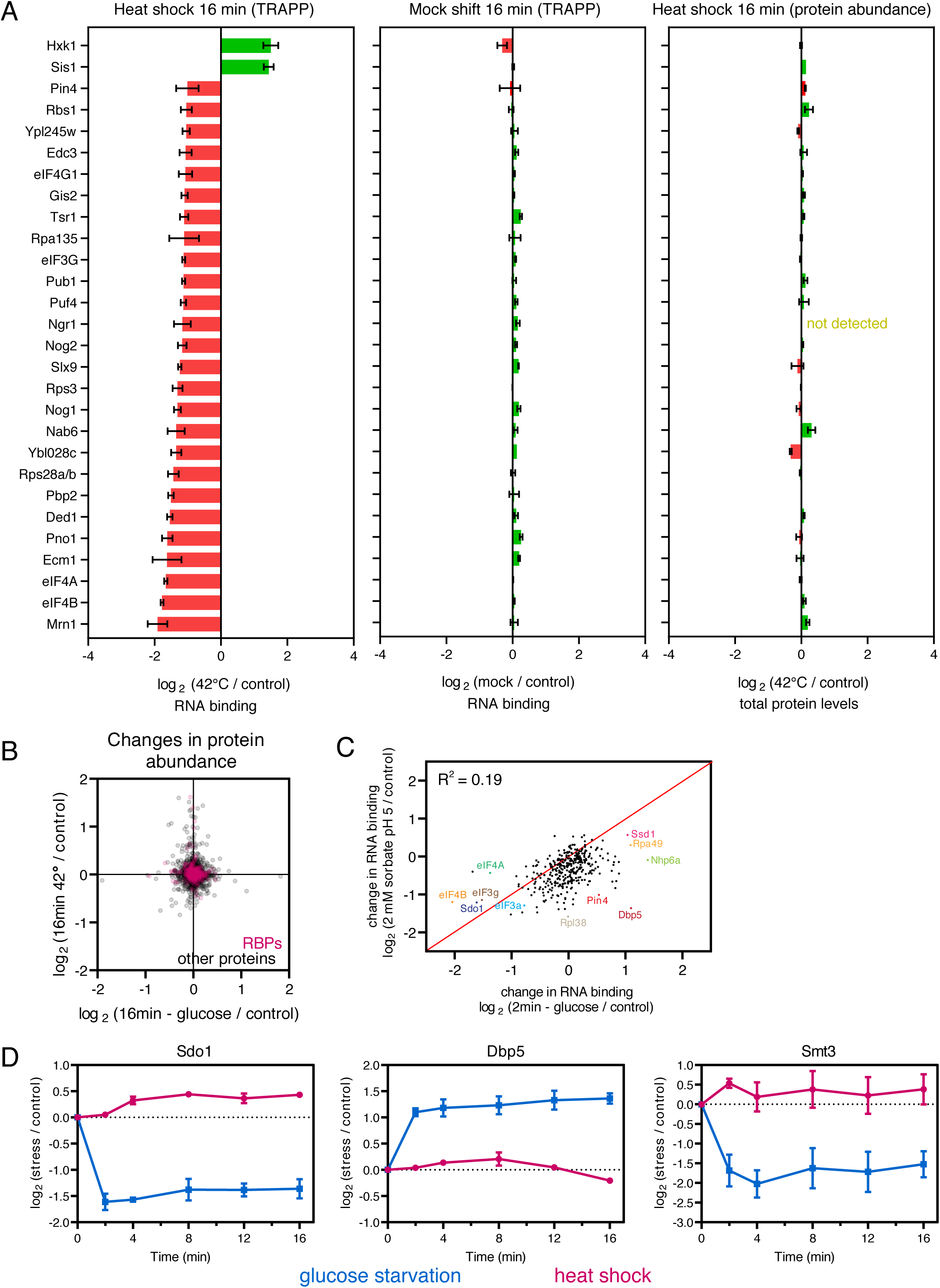
The impact of stress on the yeast RNA binding proteome. Related to Figure 1. (A) Bar chart showing all proteins with greater than 2-fold change in RNA association after 16 min of heat shock *(left)* or a mock shift *(center).* The right-hand panel shows changes in protein abundance following 16 min of heat shock *(right).* (B) Global changes in protein abundance following glucose starvation or heat shock. RNA binding proteins (RBPs) are highlighted in pink. (C) Comparison of changes in RNA association for individual RBPs after 2 min of glucose starvation, versus treatment for 2 min with 2 mM sorbate at pH 5. (D) Time course showing changes in RNA association for selected proteins.

**Figure S2.**
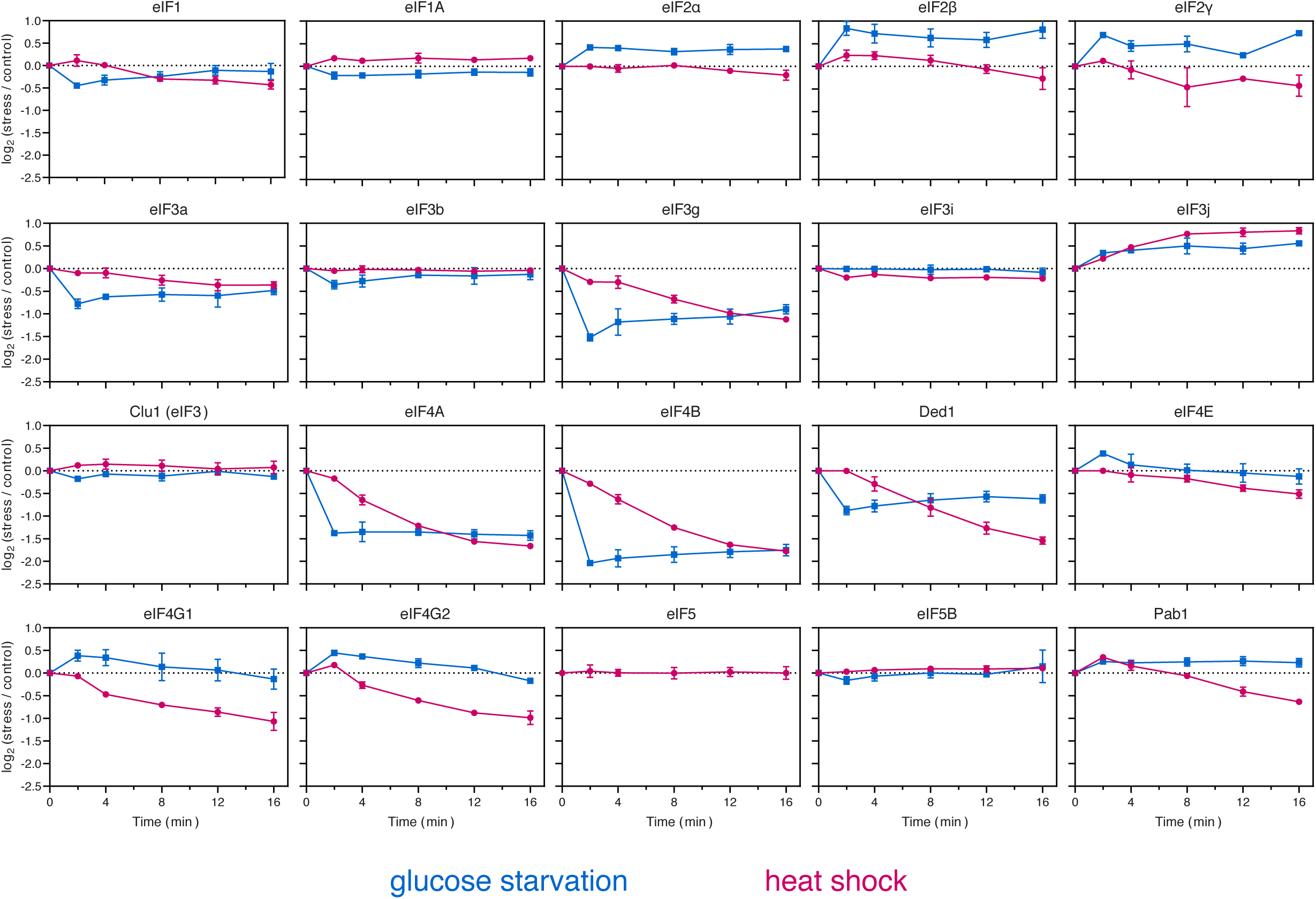
Changes in RNA binding among translation initiation factors. Related to Figure 3. Time course showing changes in RNA binding for all translation initiation factors following glucose starvation (blue) and heat shock (pink).

**Figure S3.**
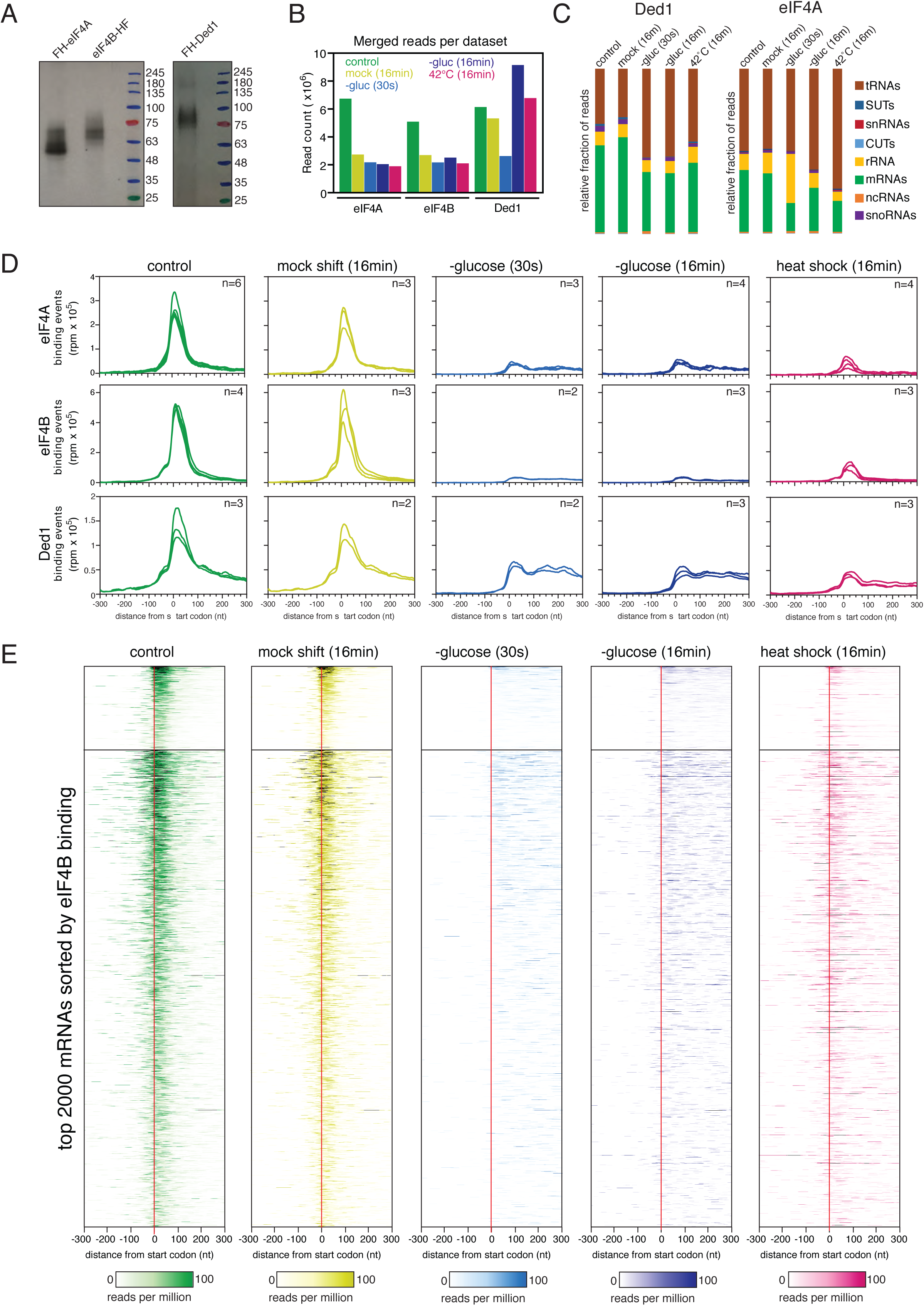
Genome-wide analysis of the RNA binding profiles of eIF4A, eIF4B, and Ded1. Related to Figure 4. (A) Autoradiogram of protein:RNA complexes separated by SDS-PAGE. (B) The total number of reads in each dataset. Reads from separate replicates were merged together. (C) Breakdown of Ded1- and eIF4A-bound RNAs by biotype. (D) Metaplots showing the distribution of eIF4A, eIF4B, and Ded1 binding around the mRNA start codon for individual replicates. The number of replicates is shown in the upper right corner. (E) Heatmaps showing the distribution of eIF4B binding around the mRNA start codon, sorted by decreasing binding. Scales are shown at the bottom. The upper part of each heatmap is scaled differently (0-1000) due to the very high number of read mapping to those transcripts. Regions that exceed the scale are colored black.

**Figure S4.**
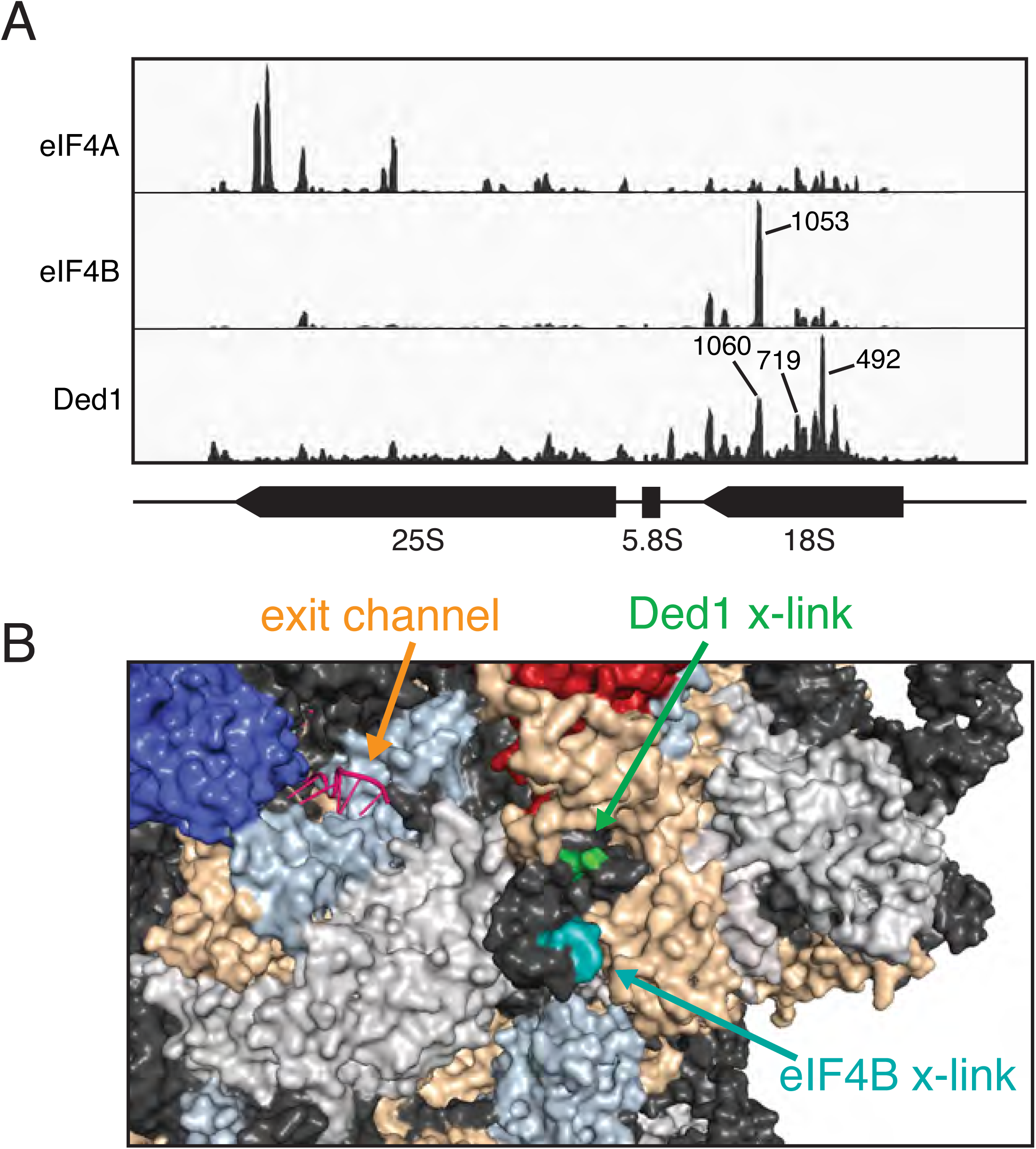
Binding of eIF4A, eIF4B, and Ded1 to ribosomal RNA. Related to Figure 4. (A) Binding of eIF4A, eIF4B, and Ded1 across the 35S pre-rRNA. (C) A close-up view of the mRNA exit channel showing the crosslinking sites on 18S for Ded1 (green) and eIF4B (teal).

**Figure S5.**
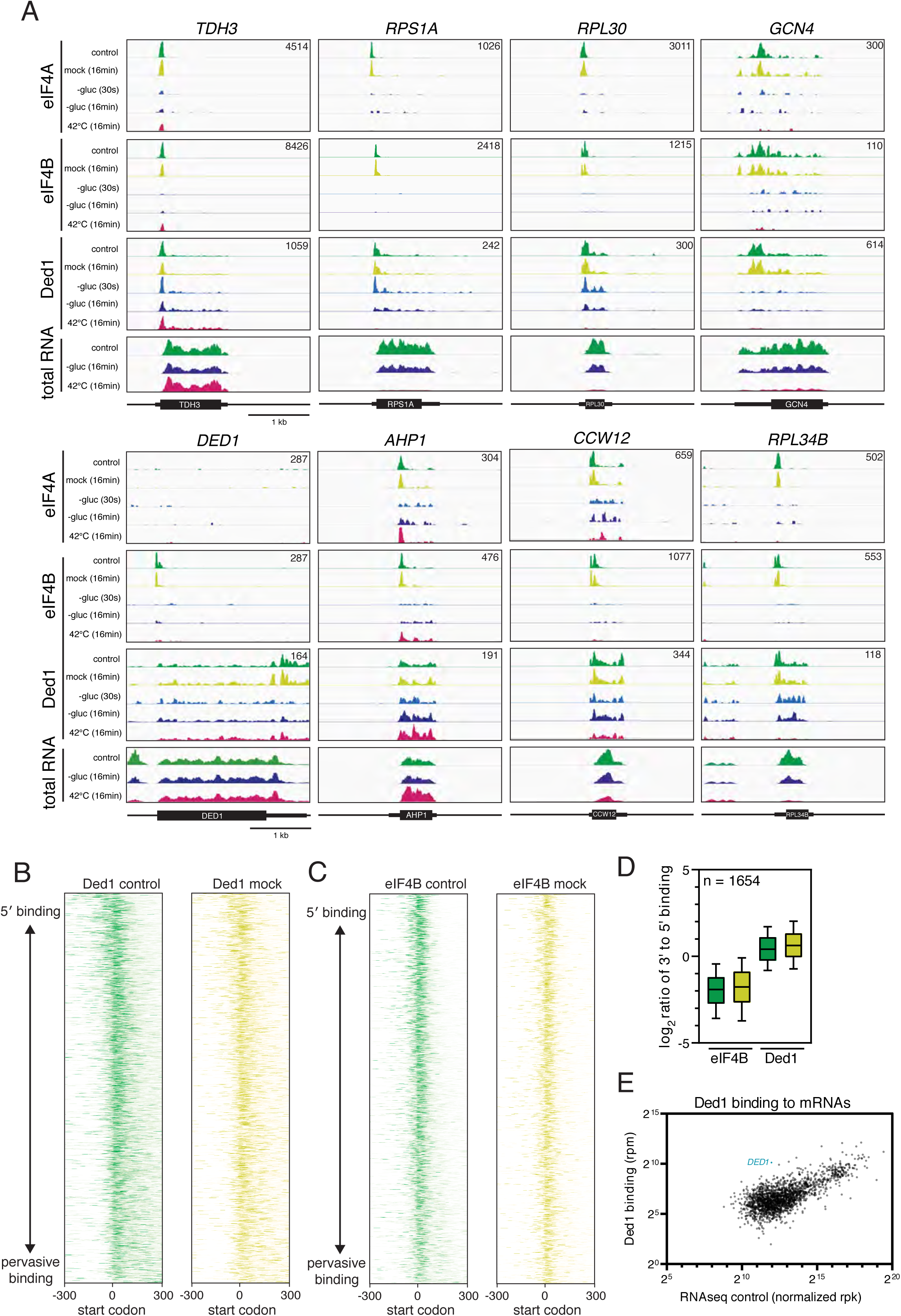
Binding of eIF4A, eIF4B, and Ded1 to specific transcripts. Related to Figure 4. (A) Binding of eIF4A, eIF4B, and Ded1 across a number of selected mRNAs. Each set of tracks is normalized to total library size using reads per million, with the exact value indicated in the upper right corner of each box. RNAseq traces are shown at the bottom as a control. Each track is normalized to a spike-in control, and thus represents the absolute abundance of each mRNA compared to the control. (B) Heatmaps showing the distribution of Ded1 around the mRNA start codon. Transcripts are sorted by the ratio of 5′ binding (5’ UTR to +150 from start codon) versus downstream binding in the control sample for each species. Only transcripts longer than 700 nt were included in the analysis (n = 1,654). (C) Same as (B) but for eIF4B. (D) Boxplot quantifying the ratio of 3’ binding to 5’ binding for eIF4B and Ded1. (E) Scatterplot comparing Ded1 binding to mRNA levels. The *DED1* transcript itself is highlighted in teal.

**Figure S6.**
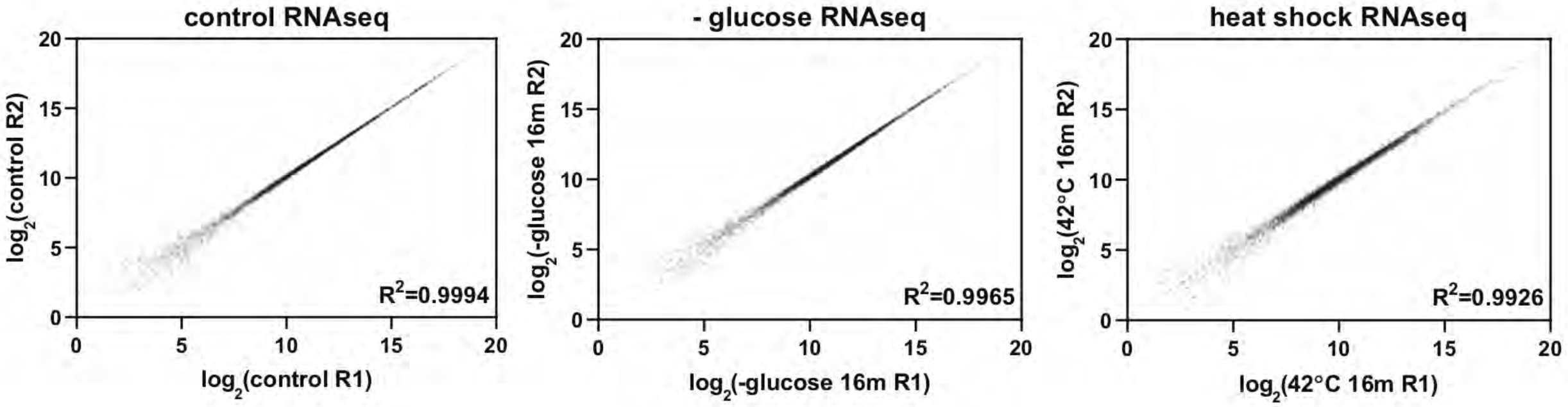
Comparison of replicate RNAseq datasets. Related to Figure 5. Scatter plots comparing RNAseq data from different replicates for all mRNAs for which sequence reads were recovered (n=6,273).

## Supplementary Table Legends

**Table S1.** Yeast strains used in this study.

| **Yeast name** | **Genotype** | **Source** |
| --- | --- | --- |
| BY4741 | MATa his3Δ1 leu2Δ0 met15Δ0 ura3Δ0 | |
| ySB076 | MATa his3Δ1 leu2Δ0 met15Δ0 ura3Δ0 lys9Δ arg4Δ | Shchepachev et al., 2019 |
| ySB082 | MATa his3Δ1 leu2Δ0 met15Δ0 ura3Δ0 xrn1Δ | haploid deletion collection |
| ySB123 | MATa his3Δ1 leu2Δ0 met15Δ0 ura3Δ0 FH-TIF1 | This study |
| ySB124 | MATa his3Δ1 leu2Δ0 met15Δ0 ura3Δ0 TIF3-HF | This study |
| ySB131 | MATa his3Δ1 leu2Δ0 met15Δ0 ura3Δ0 FH-Ded1 | This study |
| ySB151 | MATa his3Δ1 leu2Δ0 met15Δ0 ura3Δ0 ski2Δ | This study |

**Table S2.** Plasmids used in this study.

| **Plasmid name** | **Description** | **Source** |
| --- | --- | --- |
| pML104 | CRISPR-Cas9 plasmid with URA3 | Laughery et al., 2019 |
| pML107 | CRISPR-Cas9 plasmid with LEU2 | Laughery et al., 2019 |
| pSB051 | pML104-TIF3 targeting gRNA | This study |
| pSB053 | pML107-TIF1 targeting gRNA | This study |
| pSB064 | pML104-DED1 targeting gRNA | This study |
| pSB077 | pML104-SKI2 targeting gRNA | This study |

**Table S3.** DNA oligonucleotides used in this study.

**Table S4.** TRAPP results from all replicates and conditions tested. Each value indicates the log2 fold change in RNA binding during stress compared to the control. The headings for each dataset indicate the strain (e.g. wt), the stress condition, the timepoint, the replicate number, and a unique identifier associated with each sample.

**Table S5.** Changes in total protein abundance after 16 min of glucose starvation or heat shock. Each value indicates the log2 fold change in RNA binding during stress relative to the control.

**Table S6.** eIF4A CRAC results showing the amount of binding (in reads per million) to different mRNAs.

**Table S7.** eIF4B CRAC results showing the amount of binding (in reads per million) to different mRNAs.

**Table S8.** Ded1 CRAC results showing the amount of binding (in reads per million) to different mRNAs.

**Table S9.** Raw data for the heatmaps in Fig. 4E.

**Table S10.** RNAseq results from control or following 16 min of glucose starvation or heat shock. Values show reads per kilobase and were normalized to the *S. pombe* spike-in control.

## MATERIALS AND METHODS

### Cell culture and medium

All yeast strains were cultured at 30°C in synthetic medium containing 2% glucose to 0.4 OD_600_. Cells were either harvested directly (control), or collected by filtration and transferred to medium containing glucose (mock shift), medium lacking glucose but containing 2% glycerol and 2% ethanol (glucose starvation), or to glucose-containing medium prewarmed to 42°C (heat shock). Cycloheximide was used at a concentration of 0.1 mg/mL.

### Plasmid construction

All proteins were tagged using CRISPR-Cas9 as described (Laughery et al., 2015). eIF4B and Ded1 were tagged using the pML104 vector backbone (Addgene: 67638) while eIF4A was tagged using pML107 (Addgene: 67639). Both plasmids included an ampicillin resistance gene, a Cas9 expression construct, and a guide RNA (gRNA) cloning site, but differed in selectable marker (*URA3* and *LEU2*, respectively). Plasmid DNA was prepared from dam^**-**^ *E. coli.* For each plasmid, 10 µg were digested overnight with SwaI, and then for 2 h at 50°C with BclI. The digested vector was purified by gel extraction (Qiagen).

Guide RNA oligos were designed as reported (Laughery et al., 2015). Each oligo pair was annealed in a reaction consisting of 1 µM forward oligo, 1 µM reverse oligo, 50 mM Tris-HCl 7.5, 10 mM MgCl_2_, 1 mM ATP, and 10 mM DTT in a 100 µL reaction volume. The hybridization reaction was initially incubated at 95°C for 6 min, and gradually decreased to 25°C at the rate of 1.33°C/min. Hybridized substrates were then ligated into the digested vector at 25°C for 4 h. The ligation reaction consisted of 265 ng vector, 0.8 nmol insert, 50 mM Tris-HCl 7.5, 10 mM MgCl_2_, 1 mM ATP, 10 mM DTT, and 800 units of T4 DNA ligase (NEB M0202L) in a 40 µL reaction volume. The ligation mix was transformed into homemade DH5α *E. coli*, and plated overnight on LB-Amp. DNA was isolated from several colonies and sequenced to ensure correct insertion of the guide sequence.

### Strain construction

All *S. cerevisiae* strains used in this study were derived from the BY4741 background. For SILAC experiments, we used a strain auxotrophic for lysine and arginine biosynthesis (BY4741 *Δlys9 Δarg4*).

For CRAC experiments, the chromosomal copies of *TIF1* (one of two genes encoding eIF4A) and *DED1* were N-terminally tagged with FH (Flag-His), consisting of a single Flag motif, a four-alanine spacer, and eight consecutive histidine residues (DYKDDDDKAAAAHHHHHHHH). *TIF3* (eIF4B) was C-terminally tagged with the same elements in reverse (HHHHHHHHAAAADYKDDDDK), the HF (His-Flag) tag. For the *DED1* tagging, we appended an additional amino acid (asparagine) to the N-terminus of the tag to remove the gRNA cleavage site. All strains were generated using CRISPR as described below (Laughery et al., 2015).

To generate repair templates, we designed fragments consisting of the HF or FH DNA sequence flanked by 50 bp homology arms. Typically, synonymous mutations were used to disrupt the PAM site to prevent any further cleavage by Cas9. Each repair template was made by annealing two single stranded oligo nucleotides sharing 20 bp of complementarity at their 3’ ends. Each oligo pair was annealed in a reaction consisting of 10 µM forward oligo, 10 µM reverse oligo, 50 mM NaCl, 10 mM Tris-HCl 7.9, 10 mM MgCl_2_, and 100 µg/mL BSA in a 43 µL reaction volume. The hybridization reaction was initially incubated at 95°C for 6 min, and gradually decreased to 25°C at the rate of 1.33°C/min. Subsequently, the annealed oligos were incubated in the same buffer supplemented with 250 µM dNTPs and 5U Klenow exo- (NEB M0212L) in a 50µL reaction at 37°C for 1 h to fill in the single stranded regions.

To tag the genes of interest, BY4741 yeast were transformed using the standard LiOAc protocol with 500 ng of gRNA plasmid and 10 pmol of the corresponding repair template. Transformants were plated onto either leu- or ura-medium. After three days, several clones from each transformation were plated again on selective medium, and allowed to grow for an additional 2-3 days. Single colonies were selected and plated on YPD for 2 days. Finally, individual colonies were grown overnight in liquid YPD and frozen. The clones were verified by PCR using flanking primers and confirmed by sequencing. The *SKI2* gene was deleted using a guide RNA targeting the end of the open reading frame using the same approach as with the tagging.

All yeast strains used in this study are listed in Table S1. DNA oligonucleotides and plasmids used for strain construction are listed in Table S2 and S3, respectively.

### TRAPP

For TRAPP experiments, cells were cultured at 30°C in 700 mL of synthetic complete (SC) –arg -lys -trp -ura (Formedium DCS1339), supplemented with 20 µg/mL uracil (Sigma-Aldrich U0750-100G) and 2% glucose. Light media additionally included 30 µg/mL lysine (Sigma-Aldrich L5626-100G), and 5 µg/mL arginine (Sigma-Aldrich A5131-100G), while heavy media included 30 µg/mL _13_C_6_ lysine (CK Isotopes CLM-2247-H), and 5 µg/mL _13_C_6_ arginine (CK Isotopes CLM-2265-H). For most experiments, light labelling was used for cells exposed to stress, while heavy labelling was used for control cells. Prior to each experiment, cells were cultured with heavy isotope for at least 8 generations to ensure complete labelling.

Overnight starter cultures were inoculated into fresh media at a starting OD_600_ of 0.05. At OD_600_ 0.15, 4-thiouracil (4tU) (Sigma-Aldrich 440736-1G) was added to the media at a final concentration of 0.5 mM, and the cells were grown for another 200 minutes (approx. OD_600_ 0.4). After 4tU treatment for three hours, heavy labelled cells were collected by filtration and transferred to 700 mL of heavy media lacking 4tU. The UV lamps were allowed to warm up for one minute, before the shutters were opened and the cells were crosslinked for 38 sec (350 nm; 7.3 J cm^-2^). Cells were harvested by filtration, resuspended in 50 mL ice-cold PBS, and 200 ODs were collected by centrifugation. The cell pellets were frozen for later processing. Light-labelled cells were collected by filtration and quickly (3-4 sec) transferred to medium containing 2% each of glycerol and ethanol instead of glucose (glucose starvation), or standard medium pre-warmed to 42°C (heat shock). Cells for each time point were grown separately and harvesting times were sufficiently staggered to allow enough time to collect each time point. For the earliest time point, the cells were transferred to the appropriate medium lacking 4tU and crosslinked at 2 min. The cells were then processed as described above for the control sample. For all other time points, the cells were first transferred to medium containing 4tU and cultured for the appropriate amount of time. Two min prior to the end point, the cells were collected by filtration and transferred to media lacking 4tU. The cells were then crosslinked and processed as described above.

In preparation for TRAPP, 10 g of silica sand was left overnight in 50 mL of 1M HCl. The sand was then washed with 50 mL water three times (2,000 g; 2 min). After the final wash, the sand was resuspended in equivolume water to achieve a 50% slurry suspension.

Matching SILAC pairs were each resuspended in 1 mL of a 1:1 mix between phenol pH 8 and GTC lysis buffer (4 M guanidine thiocyanate, 50 mM Tris-HCl pH 8.0, 10 mM EDTA, and 1% β-mercaptoethanol), and combined in equal proportion (400 ODs total in 2 mL phenol-GTC) in a 50 mL conical. Three mL of zirconia beads was added and the cells were vortexed for 6 min to lyse the cells. An additional 8 mL of phenol-GTC was added, and the cells were vortexed for an additional 1 min. The cell lysate was centrifuged at 4,600rpm in a Sorvall centrifuge for 5 min. The supernatant was transferred to several 2 mL Eppendorf tubes, and centrifuged at 16000g for 10 min. The cleared lysate was pooled in a fresh 50 mL conical, and added to 0.1 volumes of 3 M sodium acetate pH 4.0 and mixed. An equal volume of ethanol was slowly added to the mix, followed by 1 mL of 50% silica slurry and an additional 500 µL of ethanol. The lysate was incubated at room temperature on a rotating wheel for 30 min.

The silica sand was washed three times at 2,000 rpm for 2 min with 10 mL of wash buffer I (4 M guanidine thiocyanate, 1 M sodium acetate pH 4, and 30% ethanol), followed by three washes with 10 mL of wash buffer II (100 mM NaCl, 50 mM Tris-HCl pH 6.4, and 80% ethanol). The silica was resuspended in ∼3.5 mL wash buffer II, transferred to two 2 mL Eppendorf tubes, and centrifuged at 2,000 g for 2 min. The supernatant was removed and the tubes were centrifuged in a SpeedVac for 20 min at 45°C to remove residual wash buffer.

Protein:RNA complexes were eluted three times with 10 mM Tris pH 8.0. For each elution, the silica was thoroughly resuspended in 500 µL of elution buffer, and incubated with shaking at 37°C for 5 min. The resulting eluates were combined and centrifuged at max to remove residual silica. The supernatant was removed, centrifuged a second time, and the resulting supernatant was transferred to protein LoBind tubes. The eluates were incubated with 0.25 µL of RNaseA/T1 for two hours at 37°C, followed by centrifugation overnight in a SpeedVac at room temperature.

The protein in each tube was resuspended in 35 µL of 1.5X Laemmli buffer (90 mM Tris-HCl pH 6.8, 3% SDS, 15% glycerol, and 8% β-mercaptoethanol). Matching samples were combined into a single tube, and incubated for 5 min at 100°C. Approximately 25 µL was loaded onto a 4-20% Miniprotean TGX gel and run in Tris-Glycine running buffer. Individual samples were typically split across two lanes to avoid overloading. Each gel was run at 50 V for 40 min (approximately 2 cm), and then placed in a 15 cm petri dish and rinsed with distilled water for approximately 30 min. Subsequently, the gel was stained with Imperial Protein Stain (Thermo) for 1 h, rinsed several times with water, and allowed to destain in water for 3 h to overnight.

Protein smears were cut into two sections consisting of high- and low-molecular weight proteins. Each gel fragment was diced into smaller pieces roughly 1 mm^3^ in size and collected in a 1.5 mL Eppendorf tube. The gel pieces were destained in a solution consisting of 50 mM ammonium biocarbonate (Sigma) and 50% acetonitrile for 30 min at 37°C with shaking at 750 rpm.

Proteins were then digested with trypsin as described by (Shevchenko et al., 1996). Briefly, proteins were reduced with 10 mM dithiothreitol (Sigma Aldrich, UK) in ammonium bicarbonate for 30 min at 37°C and alkylated with 55 mM iodoacetamide (Sigma Aldrich, UK) in ammonium bicarbonate for 20 min at ambient temperature in the dark. They were then digested overnight at 37°C with 13 ng/μL trypsin (Pierce, UK).

For this and subsequent steps, enough solution was added to cover the gel pieces completely. Subsequently, the gel fragments were treated with acetonitrile for 5 min to further shrink them. The acetonitrile solution was removed, and disulfide bonds were reduced with 10 mM dithiothreitol (Sigma-Aldrich) in a 50 mM ammonium bicarbonate solution for 30 min at 37°C with shaking. The DTT was removed, and the gel pieces were again shrunk by 5 min incubation with acetonitrile. Subsequently, the gel fragments were treated with 55 mM iodacetamide and 50 mM ammonium bicarbonate (Sigma-Aldrich) to alkylate free cysteines. The samples were digested overnight with trypsin in buffer consisting of 10 mM ammonium bicarbonate and 10% acetonitrile.

Following trypsin digestion, the samples were acidified to pH 1-2 using 10% trifluoroacetic acid (TFA) and processed using the stage-tip method (Rappsilber et al., 2007). Briefly, three C-18 discs were cut out and placed in a 200 µL pipet tip with gentle compression. Each stage tip was placed in a 1.5 mL collection tube with a hole in the lid to hold the stage tip. The stage tips were conditioned with washes of 40 µL methanol followed by 80 µL of 0.1% TFA. Subsequently, the peptide solution was loaded on the stage tip and centrifuged at 1000 g, to allow the peptide to bind the column. Once all of the solution had passed through, the stage tips were loaded with 25 µL of 0.1% TFA, and temporarily placed at 4°C.

In parallel, the remaining gel fragments were incubated for 10 min in a solution consisting of 80% ACN and 0.1% TFA in order to remove any remaining peptides from the gel. This solution was then transferred to a 2 mL Protein LoBind tube (Eppendorf) and dried under vacuum centrifugation at 60°C. Afterwards, the protein pellet was resuspended in 200 µL of 0.1% TFA and passed through the stage tip. Finally, the stage tip was washed twice with 100 µL of 0.1% TFA, and then placed at -20°C for storage prior to mass spectrometry.

### Mass Spectrometry

Following digestion, samples were diluted with equal volume of 0.1% Trifluoroacetic acid (TFA) (Sigma Aldrich, UK) and spun onto StageTips as described by (Rappsilber et al., 2003).

Peptides were eluted in 40 μL of 80% acetonitrile in 0.1% TFA and concentrated down to 5 μL by vacuum centrifugation (Concentrator 5301, Eppendorf, UK). The peptide sample was then prepared for LC-MS/MS analysis by diluting it to 5 μL by 0.1% TFA. MS-analyses were performed on an Orbitrap Fusion™ Lumos™ Tribrid™ mass spectrometer (Thermo Fisher Scientific, UK), coupled on-line, to Ultimate 3000 RSLCnano Systems (Dionex, Thermo Fisher Scientific). Peptides were separated on a 50 cm EASY-Spray column (Thermo Fisher Scientific, UK) assembled in an EASY-Spray source (Thermo Fisher Scientific, UK) and operated at a constant temperature of 50°C.

Mobile phase A consisted of water and 0.1% formic acid (Sigma Aldrich, UK); mobile phase B consisted of 80% acetonitrile and 0.1% formic acid. The total run time per fraction was 190 min and for protein abundance samples was 160 min per fraction. Peptides were loaded onto the column at a flow rate of 0.3 μL min^-1^ and eluted at a flow rate of 0.25 μL min^-1^ according to the following gradient: 2 to 40% buffer B in 150 min, then to 95% in 16 min. For protein abundance samples the gradient was 2 to 40% mobile phase B in 120 min and then to 05% in 16 min. In both cases, samples were subjected to mass spectrometry analysis under the same conditions. Specifically, survey scans were performed at resolution of 120,000 in the orbitrap with scan range 400-1,900 m/z and an ion target of 4.0e5. The RF lens was set to 30% and the maximum injection time to 50ms. The cycle time was set to 3 sec and dynamic exclusion to 60 sec. MS2 was performed in the Ion Trap at a rapid scan mode with ion target of 1.0E4 and HCD fragmentation with normalized collision energy of 27 (Olsen et al., 2007). The isolation window in the quadrupole was set at 1.4 Thomson and the maximum injection time was set to 35 ms. Only ions with charge between 2 and 7 were selected for MS2.

### Total proteomics

Cell culturing was performed as described above for the TRAPP experiments. Approximately 50 ODs of cells were resuspended in 200 μL of TN150 (50 mM Tris-HCl pH 7.5, 150 mM NaCl, 0.1% NP-40, 5 mM β-mercaptoethanol and a protease-inhibitor cocktail (1 tablet / 50 mL) and added to 500 mL of zirconia beads in a 1.5 mL Eppendorf tube. The cells were lysed with five one-minute pulses, with cooling on ice for one minute in between. The lysate was further diluted with 0.6 mL TN150, briefly vortexed, and combined 1:1 with 3X Laemmli buffer. The sample was incubated at 100°C for five minutes and centrifuged at 20,000g for 1 min. Approximately 10μL of supernatant was loaded onto a 4-20% Miniprotean TGX gel and run in Tris-Glycine running buffer at 100 V. Subsequently, the gel was washed with water, stained with Imperial Protein Stain (Thermo) for 1 h, rinsed several times with water, and allowed to destain in water overnight. Each lane was divided cut into six fractions, and then processed as described above for TRAPP.

### RNAseq

*S. cerevisiae* BY4741 cells were grown to 0.4 OD in SC -trp media. For control samples, the cells were collected by filtration, transferred to 50 mL ice-cold PBS, and centrifuged. Cell pellets were frozen for later use. For stress samples, cells were collected by filtration and transferred to the appropriate medium for 16 min, collected by filtration, and frozen at -80°C. Biological duplicates were collected for each condition. *Schizosaccharomyces pombe* 972H cells were harvested separately and frozen for use as a spike-in control. RNA from all samples was purified using phenol:chloroform extraction. *S. pombe* RNA was spiked into each sample at a final concentration of 2%. Libraries for RNAseq were prepared by the Wellcome Trust Clinical Research Facility at Western General Hospital (Edinburgh, UK) using poly(A) mRNA magnetic isolation (NEB #E7490) and the NEBNEXT Ultra II Directional RNA Library Prep kit (NEB #7760). The libraries were sequenced using Next-Seq with single-end, 75nt output.

### CRAC

The CRAC protocol is based on (Granneman et al., 2011), with some modifications. Most importantly, we substituted the two Protein A affinity tags and the TEV cleavage site with a single Flag tag. The histidine tag was lengthened from six residues to eight.

For each CRAC experiment, 700 mL of cells were grown in SC -trp media. At OD_600_ 0.4, the cells were UV-irradiated at 254 nm with a dose of 100 mJ/cm^2^ (4-6 sec) using the Vari-X-Link crosslinker. Following crosslinking, cells were collected by filtration and resuspended in 50 mL ice-cold PBS, and then centrifuged at 4,600g for 2 min. The cell pellets were stored at -80°C. Cell pellets were resuspended in 500 μL TN150 (50 mM Tris-HCl pH 7.5, 150 mM NaCl, 0.1% NP-40, 5 mM β-mercaptoethanol and a protease-inhibitor cocktail (1 tablet / 50 mL) and added to 1.25 mL of zirconia beads in a 50 mL conical. The cells were lysed with five one-minute pulses, with cooling on ice in between. The lysate was further diluted with 1.5 mL TN150, briefly vortexed, and centrifuged at 4,600g for 5 min. The supernatant was transferred to Eppendorf tubes and spun for an additional 20 min at 16,000g. In parallel, 100 μL of magnetic anti-Flag bead slurry was washed twice with TN150. The cleared lysate was incubated with the anti-Flag beads for two hours at 4°C, with nutating. Subsequently, the beads were washed four times with TN150 (5 min nutating at 4°C) and then incubated with 200 μL of flag peptide (100 μg / mL in TBS) at 37°C with shaking for 15 min. The eluate was transferred to a fresh tube containing 350 μL TN150 and treated with RNace-IT (0.1U, 5 min, 37 °C) to fragment protein-bound RNA. The RNase reaction was quenched by transferring the eluate to a tube containing 400 mg guanidine hydrochloride. The solution was adjusted for nickel affinity purification with the addition of 27 μL NaCl (5 M) and 3 μL imidazole (2.5 M) and added to 50 μL of washed nickel beads.

Following an overnight incubation, the nickel beads were transferred to a spin column and washed three times with 400 μL WBI (6.0 M guanidine hydrochloride, 50 mM Tris-HCl pH 7.5, 300 mM NaCl, 0.1% NP-40, 10 mM imidazole, and 5 mM β-mercaptoethanol), and then three times with 600 μL 1xPNK buffer (50mM Tris-HCl pH 7.5, 10 mM MgCl_2_, 0.5% NP-40, and 5 mM β-mercaptoethanol). Subsequent reactions (80 μL total volume for each) were performed in the columns, and afterward washed once with WBI and three times with 1xPNK buffer:

1. Phosphatase treatment (1x PNK buffer, 8 U TSAP (Promega), 80 U RNasIN (Promega); 37°C for 30 min).
2. 3′ linker ligation (1x PNK buffer, 20 U T4 RNA ligase I (NEB), 20 U T4 RNA Ligase II truncated K227R, 80 U RNasIN, 1 μM preadenylated 3′ miRCat-33 linker (IDT); 25°C for 6 h).
3. 5′ end phosphorylation and radiolabeling (1x PNK buffer, 40 U T4 PNK (NEB), 40 μCi _32_P-γATP; 37°C for 60 min, with addition of 100 nmol of ATP after 40 min).
4. 5′ linker ligation (1x PNK buffer, 40 U T4 RNA ligase I (NEB), 80 U RNasIN, linker, 1 mM ATP; 16°C overnight).

The beads were washed twice with WBII (50 mM Tris-HCl pH 7.5, 50 mM NaCl, 0.1% NP-40, 10 mM imidazole, and 5 mM β-mercaptoethanol). Protein:RNA complexes were eluted twice (10 minutes each) in 40 μL of elution buffer (same as WBII but with 300 mM imidazole). At this point, different replicates or conditions for the same protein were combined. The merged eluates were precipitated with 5X volume acetone at -20°C for at least two hours. RNPs were pelleted at 16000g for 20 min, and resuspended in 20 μL 1X NuPAGE sample loading buffer supplemented with 8% β-mercaptoethanol. The sample was denatured by incubation at 65°C for 10 min, and run on a 4%–12% Bis-tris NuPAGE gel at 150 V. The protein:RNA complexes were transferred to Hybond-C nitrocellulose membranes with NuPAGE MOPS transfer buffer for 1.5 h at 100V. Labelled RNA was detected by autoradiography. The appropriate region was excised from the membrane and treated with 0.25 μg/μL Proteinase K (50 mM Tris-HCl pH 7.5, 50 mM NaCl, 0.1% NP-40, 10 mM imidazole, 1% SDS, 5 mM EDTA, and 5 mM β-mercaptoethanol; 2 hr 55°C with shaking) in a 500 μL reaction. The RNA component was isolated with a standard phenol:chloroform extraction followed by ethanol precipitation. The RNA was reverse transcribed using Superscript III and the miRCat-33 RT oligo (IDT) for 1 hr at 50°C in a 20μL reaction. The resulting cDNA was amplified by PCR in five separate reactions using La Taq (2 μL template, 18-21 cycles) PCR reactions were combined, precipitated in ethanol, and resolved on a 3% Metaphore agarose gel. A region corresponding to 140 to 200 bp was excised from the gel and extracted using the Min-elute kit. Libraries were sequenced by the Wellcome Trust Clinical Research Facility (Edinburgh, UK) on Next-Seq with single-end, 75nt output.

### RT-qPCR

Approximately 10 mL of cells at OD 0.4 were collected and added to 40 mL of ice-cold medium to rapidly cool the cells. The cells were pelleted by brief centrifugation (4,600 rpm; 1 min), resuspended in 1mL of ice cold PBS, centrifuged again to pellet, and frozen at -80°C. Total RNA was isolated by the GTC-phenol method. Residual DNA was degraded by treating with Turbo DNase (Invitrogen) in a 50 μL reaction consisting of 2 µg of RNA, 1 U DNase and 20 U of RNasin in a 1X solution of the manufacturer-supplied buffer at 37°C for 30 min. DNase was removed by phenol-chloroform-isoamyl alcohol extraction followed by ethanol precipitation. Glycoblue (Life Technologies) was used as a coprecipitant and to visualize the RNA pellet after centrifugation. cDNA was synthesized with SSIII reverse transcriptase (Life Technologies) using 200 ng random hexamers (Thermo Scientific) according to the manufacturer’s protocol. Expression levels of individual transcripts were determined by quantitative PCR using SYBR Green (Roche) for detection. cDNA dilutions of 1:5 were used for amplifying mRNAs, while a dilution of 1:400 was used for amplifying rRNA. Each sample was measured in technical triplicate and the results were averaged together. Relative expression was calculated using ΔΔC_t_ and by normalizing to the levels of 25S rRNA in each sample.

## DATA ANALYSIS

### Mass spectrometry analysis

The MaxQuant software platform (Cox and Mann, 2008) version 1.6.1.0 was used to process the raw files and search was conducted against *Saccharomyces cerevisiae* complete/reference proteome set of UniProt database (released on 14/06/2019), using the Andromeda search engine (Cox et al., 2011). For the first search, peptide tolerance was set to 20 ppm while for the main search peptide tolerance was set to 4.5 pm. Isotope mass tolerance was 2 ppm and maximum charge to 7. Digestion mode was set to specific with trypsin allowing maximum of two missed cleavages. Carbamidomethylation of cysteine was set as fixed modification. Oxidation of methionine and acetylation of the N-terminal were set as variable modifications. Multiplicity was set to 2 and for heavy labels Arginine 6 and Lysine 6 were selected. Peptide and protein identifications were filtered to 1% FDR. Only proteins identified with high confidence (peptides ≥2) were considered for further analysis.

### PCA

For the PCA analysis in Fig. 1F, all TRAPP datasets were included. The raw log2 values were mainly between -2 and +2, and data normalization was not required. For each protein, the median value between replicates was used. In total, 302 proteins were present in at least two biological replicates for each timepoint and condition, and thus included in the analysis. For the PCA in Fig. 1G, only the 16 min timepoints from glucose starvation, heat shock, and mock shift were included. PCA was calculated with python v3.6.8 using the Jupiter notebook and the scikit-learn v0.22.2 library.

### Ribosome structure

The ribosome structure 3J77 (Fig. 2) is derived from (Svidritskiy et al., 2014). The mRNA was modelled into the structure, using 3J81(Hussain et al., 2014).

### CRAC analysis

The datasets were dumultiplexed using pyBarcodeFilter from the pyCRAC package (Webb et al., 2014). Flexbar (Dodt et al., 2012) was used to remove sequencing adapters, trim low-quality positions from the 3′ end, and remove low-quality reads (parameters: -ao 4 -u 2 -q TAIL -m 11 - at RIGHT with adapter sequence TGGAATTCTCGGGTGCCAAGGC). In addition to the barcode, each read contained three random nucleotides at the 5′ end to allow PCR duplicates to be removed by collapsing identical sequences with pyFastqDuplicateRemover (Webb et al., 2014). Reads were filtered to exclude low-entropy sequences using bbduk (sourceforge.net/projects/bbmap/) with parameters entropy=0.5 entropywindow=10 entropyk=6. Because translation initiation factors largely target spliced mRNAs, the sequencing reads were mapped to a modified version of the *S. cerevisiae* EF4.74 genome (Ensembl) in which the introns had been bioinformatically removed. The reads were aligned using Novoalign, with reads mapping to multiple locations randomly assigned (-r Random). The numbers of reads mapping to different mRNAs were determined using pyReadCounters (Webb et al., 2014) and a custom genome annotation file. Binding to individual mRNAs was quantified in one of two ways: 1) ‘total binding’, reflecting the number of reads mapping anywhere within a given transcript, and 2) ‘5′ end binding’, reflecting the number of reads mapping within the 5’ UTR and the first 150 nt of coding sequence. Three pseudocounts were added to each transcript to improve the quantification. Most analyses (Fig. 4B-E, S4D-E) were based on the list of 2,000 transcripts showing the strongest binding to eIF4B, defined by the reads per million average between the ‘control’ and ‘mock shift’ conditions. In order to visualize binding across individual transcripts (e.g. Fig. 4F), the coverage at each position along the genome was calculated and normalized to the library size using genomecov from bedtools. The Integrative Genomics Browser was used to visualize binding across individual transcripts (Robinson et al., 2011). Coverage around start codons (Fig. 4E and S4D-E) was calculated using pyBinCollector (Webb et al., 2014). The resulting metaplots were generated in GraphPad Prism 8, while the heatmaps were made using Excel.

### RNAseq data analysis

RNAseq reads were aligned to a concatenated genome consisting of the intronless *S. cerevisiae* genome and the 972H *Schizosaccharomyces pombe* genome. The reads mapping to the *S. pombe* genome were tabulated using pyReadCounters together with a custom annotation file. These values were used as a normalization standard to allow for quantitative comparisons between datasets. Genome coverage files (e.g. Fig. 4F) were generated using genomecov from bedtools and scaled using values determined from the *S. pombe* spike-in control. For all analyses, we used the top 5,000 transcripts, defined by their average expression across all three conditions (control, glucose withdrawal 16 min, and heat shock 16 min). GO analysis was performed using GOrilla (Eden et al., 2009) on the 500 genes showing the greatest decline in mRNA levels. Violin plots were generated using GraphPad Prism 8.

## Notes

### Competing Interest Statement

The authors have declared no competing interest.

